# Folding-Driven Control of the Functional PP1 Complex via Multiscale Modeling

**DOI:** 10.64898/2026.09.23.753607

**Authors:** Federico Fontana, Bryan Bogin, Michael Irvin, Mustafa Ozen, Zachary A Levine, Carlos F. Lopez

**Author notes:** **Corresponding Author: Carlos F Lopez** – Bay Area Institute of Science, Altos Labs Inc. - **Email**. Unità di Ingegneria Tissutale, Fondazione IRCCS Casa Sollievo della Sofferenza, Viale Cappuccini 1, San Giovanni Rotondo, Foggia 71013, Italy (Federico Fontana).

## Abstract

Understanding how conformational dynamics regulate protein complex assembly remains a central challenge in molecular systems biology. Here, we develop a multiscale modeling framework that integrates all-atom molecular dynamics (MD) simulations with rule-based kinetic modeling to investigate formation of the PP1–GADD34–eIF2α complex, a critical regulator of the integrated stress response. Using umbrella sampling, we compute the potential of mean force (PMF) for key interactions, revealing strong thermodynamic driving forces for eIF2α binding (ΔG = –89.5 kJ/mol, KD ≈ 0.07 fM) and moderate affinity for PP1–GADD34 association (ΔG = –40.7 kJ/mol, KD ≈0.138 μM). These energetics inform a PySB-based model that incorporates GADD34’s folding state as a continuous variable (ϕ), linking conformational transitions to holoenzyme assembly and activity. Simulations show that catalytic efficiency is maximized when GADD34 folding free energy is 20 kJ/mol and ϕ = 0.9, reflecting a highly ordered state stabilized by actin and PP1 binding. Comparative analysis of energy-constrained and optimized parameter regimes reveals that folding-dependent assembly enhances both eIF2α dephosphorylation and information transmission, with channel capacities increasing from <0.002 bits to >5.5 bits. These results demonstrate how conformational regulation encodes system-level function and underscore the utility of multiscale models in bridging molecular energetics with dynamic biochemical control.

## INTRODUCTION

Cellular processes depend on the precise assembly of molecular complexes, often as pivotal decision points in signal transduction and network regulation.^1–3^ These assemblies are not random but are driven by highly orchestrated interactions among components, forming functional macromolecular structures.^4,5,6^ The pathways and mechanisms underlying the assembly of these complexes are inherently multiscale. They comprise atomic-level interactions that drive local binding events and mesoscale processes that govern the emergence of network-wide functionality. Despite their significance, understanding these processes remains a considerable challenge, as it requires the integration of structural, thermodynamic, and kinetic insights across spatial and temporal scales.

Molecular dynamics (MD) simulations have proven invaluable for probing protein-protein interactions at the atomic scale. By revealing atomic-level binding modes, interaction energies, and conformational changes, they provide essential details about the physical basis of molecular recognition.^7–9^ However, these simulations alone are typically constrained by timescales, limiting their ability to capture the complex assembly process, particularly for multicomponent assembly. To address this limitation, mesoscale approaches such as Chemical Reaction Networks (CRNs) offer a complementary perspective. CRNs model the interactions of molecular components as networked reactions, enabling the prediction of complex assembly pathways and rates informed by kinetic and thermodynamic data.^10–12^ Together, these approaches can potentially unravel the intricate interplay of forces driving molecular assembly.

This work presents a multiscale modeling framework that integrates atomic-scale and mesoscale methods to investigate the assembly dynamics of protein complexes. We focus on assembling the PP1-GADD34-eIF2α complex, a key regulator in the termination of the cellular stress response pathway.^13,14^ Dysregulation of this pathway is implicated in various diseases, including neurodegenerative disorders and cancer, underscoring the importance of understanding the molecular basis of its assembly. We estimate protein-protein interaction energies by employing MD simulations with Umbrella Sampling and derive kinetic parameters for individual binding events.^15,16^ These parameters are subsequently used to inform CRNs, enabling us to model the dynamics of the entire assembly process.

Our approach provides a unified framework for understanding complex assembly pathways, bridging the gap between atomic interactions and network behavior. It shows how molecular components and their interactions give rise to emergent properties and may inform therapeutic targeting of dysregulated assemblies. This work demonstrates the value of multiscale modeling in systems biology and establishes a blueprint for studying other complex assemblies in diverse cellular contexts.

## MATERIAL AND METHODS

### MOLECULAR DYNAMICS SIMULATIONS

#### STRUCTURAL PREPARATION

The initial structural model for the pre-dephosphorylation complex of phosphorylated eIF2α with the trapped holophosphatase was obtained from the Protein Data Bank (PDB; ID: 7NZM).^17^ Using the Protein Preparation Workflow in Schrödinger,^18–19^ missing atoms were added to optimize bond orders, formal charges, and hydrogen placements. Residual steric clashes were resolved using ISOLDE (https://tristanic.github.io/isolde/),^20^ which enables interactive corrections. N-terminal and C-terminal caps were added using the pdb2gmx utility in GROMACS with the CHARMM force field, and the protonation states were assigned based on physiological pH. Specifically, buried histidine, as well as arginine and lysine, were considered positively charged, whereas aspartic acid and glutamic acid were treated as negatively charged.^1,2^ In addition, the side chain of Ser52 of eIF2α has been considered as a phosphorylated residue.

#### SYSTEM SOLVATION AND NEUTRALIZATION

The prepared complex was solvated in a cubic box containing 134,000 explicit TIP3P water molecules, ensuring a minimum distance of 10 Å between the complex and the box edges. Sodium and chloride ions were added using the genion utility in GROMACS that replaces solvent molecules with ions, to neutralize the system and achieve an ionic strength of 0.150 M NaCl.

#### EQUILIBRATION

The solvated system was subjected to energy minimization using the steepest descent algorithm to resolve steric clashes. Equilibration was performed under NVT and NPT conditions (constant pressure and temperature at 310 K and 1 bar, using a v-rescale thermostat and c-rescale barostat, respectively) with position restraints, using force constants equal to 1000 kJ mol^-1^ nm^-2^, applied to heavy atoms to stabilize the system. The equilibration run was conducted with a timestep of 0.002 fs for 2,500,000 steps, corresponding to a simulation time of 5.0 ns.

#### DISTANCE RESTRAINTS AND SHORT MD SIMULATIONS

After equilibration, distance restraints were generated. Subsequently, a short MD simulation was performed under NPT conditions, constant pressure and temperature at 310 K and 1 bar, using a v-rescale thermostat and c-rescale barostat, to stabilize the system further with these restraints applied. This simulation was conducted for 500,000 steps with a timestep of 0.002 fs, resulting in a total simulation time of 1.0 ns.

#### PULLING SIMULATIONS FOR p-eif2α DISSOCIATION

To initiate the dissociation of eIF2α from the PP1 holoenzyme complex, a pulling simulation was performed using GROMACS. The simulation was conducted for 7,500,000 steps with a timestep of 0.002 fs, resulting in a total simulation time of 15.0 ns. A constant pulling speed of 0.0002 nm/ps was applied along the reaction coordinate, corresponding to a total displacement of 3.0 nm. This simulation generated the initial configurations for umbrella sampling.

#### PUSHING SIMULATION FOR p-eif2α

Following the pulling simulation, a pushing simulation was performed to confirm that the pulled configurations corresponded to a minimum-energy conformation along the reaction coordinate. This simulation was conducted for 2,500,000 steps with a timestep of 0.002 fs, resulting in a total simulation time of 5.0 ns. A constant pushing speed of 0.0002 nm/ps was applied, ensuring a careful assessment of the energy landscape.

#### UMBRELLA SAMPLING

Umbrella sampling simulations were carried out to analyze the dissociation process further, using the minimum conformation of the PP1 holoenzyme assembly as a reference frame.^3,4^ For the pulling simulation, windows were spaced at 0.05 nm intervals over a reaction coordinate range of 3.7 nm, resulting in 74 frames. For the pushing simulation, windows were spaced at 0.05 nm intervals over a range of 0.9 nm, resulting in 18 frames. Each window was equilibrated for 5 ns with position restraints before performing production simulations for 10 ns without restraints. The Weighted Histogram Analysis Method (WHAM) was employed to reconstruct the potential of mean force (PMF) profile, providing quantitative estimates of the binding and dissociation-free energy landscapes. To account for statistical noise and ensure the robustness of the results, bootstrapping techniques were applied within the WHAM framework. This involved resampling the input histograms to generate multiple replicas of the dataset, allowing the estimation of statistical uncertainties associated with the PMF. The resulting error bars reflect the variability across these resampled datasets and provide confidence intervals for the computed free energy differences. In addition, convergence of the PMF profiles was monitored by comparing independent simulation windows and ensuring sufficient overlap of histograms across reaction coordinates, thereby minimizing artifacts due to under sampling or poor equilibration.

##### PP1 dissociation from the GADD34-a dimer

Following the dissociation of eIF2α, PP1 was pulled from the GADD34-actin dimer for a similar dissociation study.

- **System Solvation and Equilibration:** The GADD34:Actin: PP1 system was solvated in a cubic box containing 214,000 TIP3P water molecules and counterions to achieve an ionic strength of 0.150 M NaCl. The system underwent energy minimization and NVT equilibration for 5 ns under position restraints, followed by 5 ns of NPT equilibration at 310 K and 1 bar.
- **Pulling Simulation:** The dissociation of PP1 from GADD34-actin was initiated using a pulling simulation conducted for 5,000,000 steps with a timestep of 0.002 fs, corresponding to a total simulation time of 10.0 ns. A constant pulling speed of 0.001 nm/ps was applied along the reaction coordinate, resulting in a total displacement of 10.0 nm.
- **Pushing Simulation:** Following the pulling simulation, a pushing simulation was performed to confirm that the system was near a minimum-energy conformation. The pushing simulation followed the same protocol as the eIF2α simulation, running for 2,500,000 steps with a timestep of 0.002 fs, corresponding to 5.0 ns.
- **PMF Calculation:** The PMF for PP1 dissociation was derived using umbrella sampling. Windows were spaced at 0.1 nm intervals along the reaction coordinate, with 100 frames for the pulling simulation and 20 for the pushing simulation. Each window was equilibrated for 5 ns with position restraints, followed by 10 ns of production simulation. WHAM was used to reconstruct the PMF profile, providing quantitative estimates of the free energy landscape for PP1 dissociation.

#### UMBRELLA SAMPLING ANALYSIS

For both dissociation studies, umbrella sampling was performed using the same procedure. The Weighted Histogram Analysis Method (WHAM) was employed to reconstruct the potential of mean force (PMF) profile, providing insights into the binding and dissociation-free energy landscapes.

#### Model of GADD34 folding and binding

The connection between the kinetic parameters and the PMF, associated with the GADD34 unfolding process, is made through the relationship between equilibrium constants and free energy, namely:

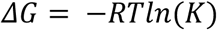

Where *K* denotes the equilibrium constant, defined as:

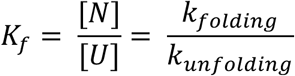

Where N is the folded state, while U denotes the unfolded state, which gives the free energy change for folding as

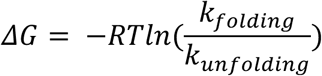

For the binding equilibrium of a ligand (L) to the folded protein (N), the binding equilibrium constant is:

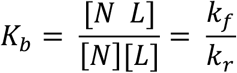

The associated free energy change for binding is:

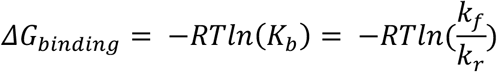

To capture how GADD34 folding affects the dynamics of complex assembly, we introduced the parameter ϕ, defined as follows:

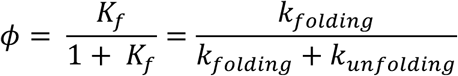

Thus, the overall observed binding constant is:

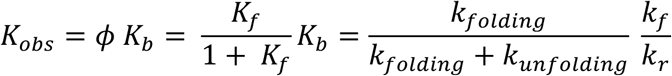

And the corresponding free energy becomes:

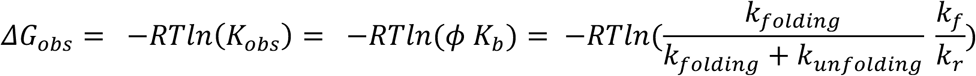

That can also be rewritten as:

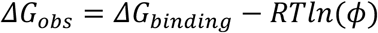

This parameter ranges from 0 to 1, where 0 indicates a fully unfolded state, while 1 represents a fully folded state. Intermediate values correspond to partial folding. Specifically:

- When *ϕ* = 1, *ΔG_obs_* = *ΔG_bindin_*_g_;
- When *ϕ* < 1the term −*RTln*(*ϕ*) becomes positive making *ΔG_obs_* less favorable. This reflects the penalty due to partial folding

##### Model of PP1 holoenzyme catalysis dependent on GADD34 folding

To capture the effect of folding on catalytic efficiency, the folding state modulates the effective energy barrier via the parameter ϕ. This implies that only the enzymes’ folded (active) fraction contributes to catalysis. In energy terms, if the intrinsic catalytic barrier is given by *ΔG_cat_*, then only a fraction ϕ of the enzyme is active. Then, this can be incorporated by writing an effective energy:

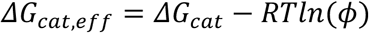

So, when ϕ=1, the catalytic barrier is *ΔG_cat_*, and if ϕ <1, the barrier is effectively higher since −*RTln*(*φ*) is positive.

#### PySB MODELING

The PP1-GADD34-eIF2α complex consists of four distinct monomeric species whose specific interactions lead to two assembly pathways. These pathways are critically dependent on the folding state of GADD34, as shown in **Figure 1**. To capture how GADD34 folding affects the dynamics of complex assembly, we introduced the parameter **ϕ**, as mentioned above. This parameter ranges from 0 to 1, where 0 indicates a completely unfolded state, 1 represents a fully folded state, and intermediate values correspond to partial folding. Essentially, ϕ describes the barycentric distance of the GADD34 population along the folding continuum.

**Figure 1.**
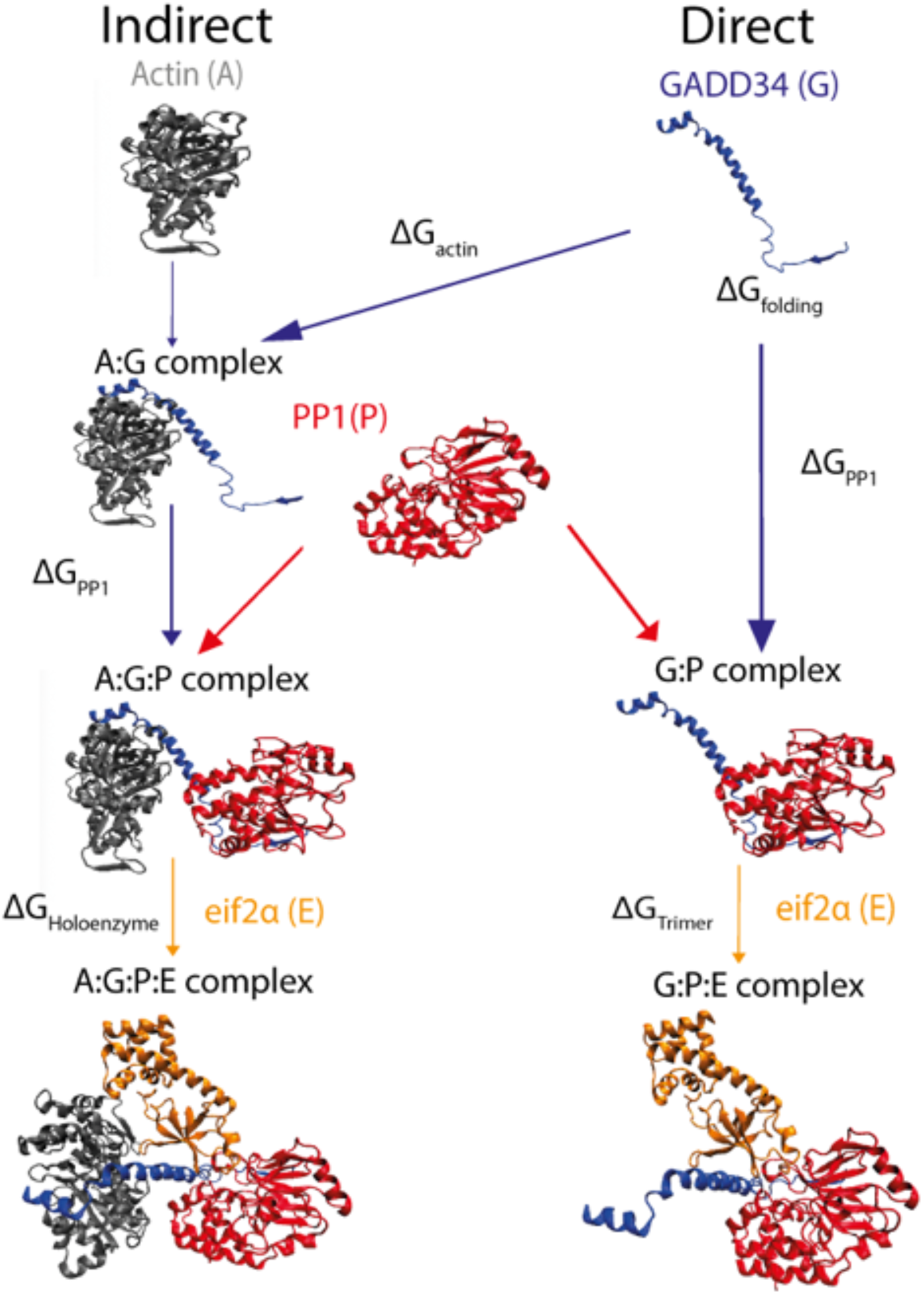
Stepwise assembly pathways of the PP1-GADD34-eIF2α complex, highlighting the molecular interactions and assembly sequence. In the indirect pathway, GADD34 (G) binds actin (A), followed by the recruitment of PP1(P) and eIF2α (E) to form the final A:G:P:E complex. On the other hand, in the direct pathway, PP1(P) might bind GADD34 (G) first, followed by the addition of eIF2α (E) and resulting in the formation of G:P:E complex.

Simultaneously, we implemented a model for forming the PP1-GADD34-eIF2B complex using PySB 1.16.0. This model encompasses 11 distinct biochemical species and 18 chemical reactions, including eight energy-constrained and two non-energy-constrained, as highlighted in **Table S1**. The model’s combinatorial complexity is evidenced by its 12 parameters, some of which were fixed based on experimental measurements from previous studies.^12^ Initial conditions were established to replicate the experimental setup accurately. More in detail, the model consists of 10 rules, 6 of them energy-based, as shown in **Figure 4**.

#### FITNESS LANDSCAPE OF HOLOENZYME PP1 ACTIVITIES

In the model, the fitness landscape is defined by the ratio:

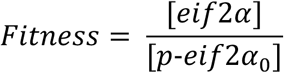

Which serves as a proxy for the efficiency of the PP1–GADD34–mediated dephosphorylation reaction. In this framework, the catalytic activity is modulated by two key parameters: the folding state of GADD34, denoted as *ϕ*(ranging from 0 to 1), and its folding free energy *ΔG_fold_*, as mentioned above.

##### Methodological Framework for Mutual Information Analysis in GADD34 Folding

To elucidate the influence of GADD34 folding on holoenzyme formation and subsequent eIF2α dephosphorylation, the mutual information for each reaction was calculated, as shown below:

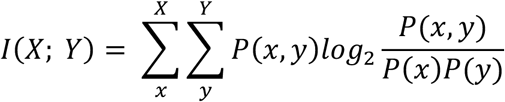

Mutual information quantifies the extent to which one variable reduces uncertainty about another. However, this calculation is challenging in systems biology because the underlying input distribution is generally unknown. The concept of channel capacity—the maximum information transmission achievable across all possible input distributions, as shown below—addresses this issue. The channel capacity framework was applied to two distinct simulation systems.

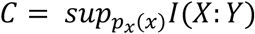

In these simulations, φ_fold was varied systematically and sampled discretely within a biologically relevant range to capture the system’s dynamic response without reliance on a fixed reference steady state. Mutual information between molecular species was estimated using the PyInform library, which applies a discrete estimator based on frequency counts of min–max normalized time series. No kernel density estimation or continuous approximation was employed. Channel capacity was approximated using a brute-force method by computing mutual information across the sampled φ values and selecting the maximum observed value to estimate maximum achievable information transfer under the tested conditions.

## RESULTS

### Folding Pathways and Allosteric Control in the PP1–GADD34–eIF2α Complex

Protein phosphatase 1 (PP1) is an important regulator of pathways, such as glycogen metabolism, muscle contraction, cell progression, neuronal activities, cell division, apoptosis, and protein synthesis.^21,22^ During transient protein synthesis inhibition, mediated by the phosphorylation of the α subunit of eukaryotic translation initiation factor 2 (eIF2α), the mRNAs encoding ATF4 and CHOP transcription factors are preferentially translated.^12,13^ Together, these factors induce the expression of the stress response gene that encodes Growth Arrest and DNA-damaged protein 34 (GADD34).^12^ As shown in **Figure 1**, GADD34 can follow different folding pathways, named Direct and Indirect, that determine its binding interactions once translated.

In one route, named Direct, depending on its folding state, GADD34 may preferentially bind directly to PP1 through its C-terminal domain, where the RVxF (555KVR558) motif residues are located, as per structural insights, such as NMR spectroscopy, X-ray crystallography investigations.^23^ Previous studies have shown that this motif is intrinsically disordered without PP1, meaning that GADD34 lacks a fixed structure until it engages with its binding partner.^13,14^ As observed with other intrinsically disordered proteins (IDPs), the folding stability of GADD34 plays a critical role in its binding dynamics.^24^

The Indirect pathway involves the binding of GADD34 to Actin, stabilizing its secondary structures and reducing the free energy barrier associated with its folding. ^25,26^ This stabilized conformation (A:G complex) favors subsequent PP1 binding and may have significant implications for the enzymatic activity of the resulting holoenzyme complex. Specifically, the proper folding of GADD34 can serve as an allosteric modulator of PP1’s catalytic activity. When GADD34 is more ordered—either through direct binding-induced folding upon association with PP1 or indirectly via actin stabilization—it may optimize the spatial orientation of PP1’s active site.^25,26^ This reorganization can enhance the recruitment and precise positioning of the substrate, eIF2α, thereby facilitating a more efficient dephosphorylation reaction.^25,26^

Conversely, if GADD34 remains partially disordered, its less defined conformation might result in suboptimal alignment of PP1’s catalytic residues with eIF2α, potentially leading to decreased catalytic turnover or altered substrate specificity. Thus, the dynamic interplay between GADD34’s folding state and its binding partners can finely tune the enzymatic output of the holoenzyme complex. Briefly, the conformational states of GADD34 govern the holoenzyme assembly route and modulate the catalytic efficiency and overall functionality of the PP1–GADD34–eIF2α complex during the stress response.

### Unraveling Molecular Recognition in PP1–GADD34–eIF2α Assembly: An Umbrella Sampling Approach

Umbrella sampling simulations were performed to unravel the molecular mechanisms underlying molecular recognition and binding reactions that lead to the assembly of PP1–GADD34–eIF2α.

The binding of **eIF2α** to the A:G:P complex, shown in **Figure 2a**, exhibits a strong thermodynamic driving force (**ΔG_on_ = -89.53 kJ/mol**) as reflected in the PMF profile. The relatively low dissociation barrier (**ΔG_D_ = 10.58 kJ/mol**) indicates that while the interaction is stable, the complex retains dynamic properties that allow reversible interactions under cellular conditions. The computed dissociation constant (**K_D_ = 0.07 fM)** confirms the strong affinity of eIF2α for the pre-assembled complex, suggesting its critical role in regulating the final steps of the assembly process. The comparative analysis of PP1 and actin recognition by eIF2α (**Figure 2c**) revealed thermodynamic preferences for PP1 binding, with **ΔG _eIF2α:PP1_ = -50.189 kJ/mol** and **K_D_ = 3.42 nM**, compared to actin binding (**ΔG _eIF2α:a_ = - 47.39 kJ/mol, K_D_ = 10.01 nM**). These differences underscore the specificity of **eIF2α** toward **PP1** in driving the assembly of the functional complex.

**Figure 2.**
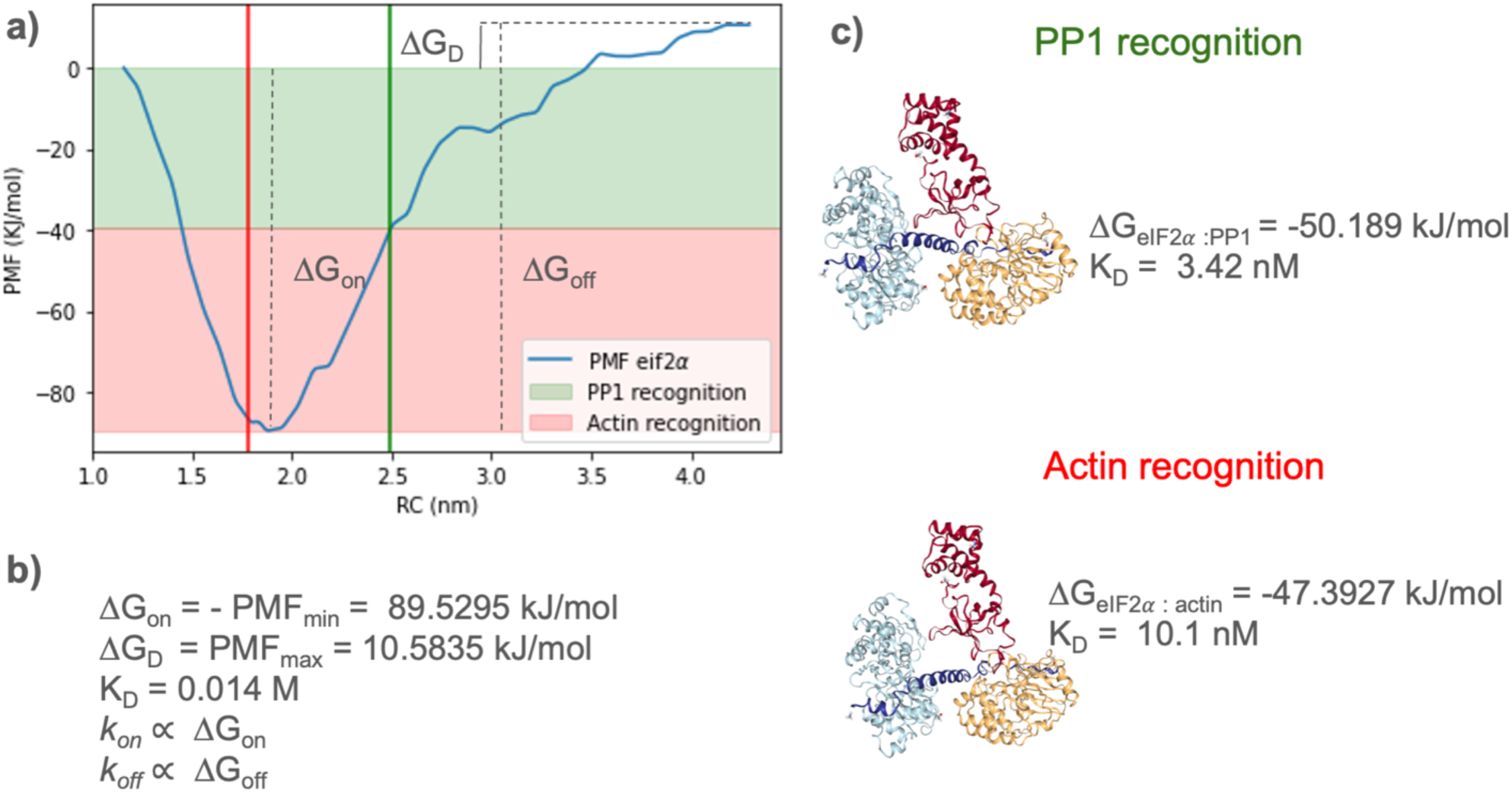
Multiscale analysis of binding interactions and recognition steps during the PP1-GADD34-eIF2α assembly. **(a)** Potential of Mean Force (PMF) profile for eIF2α binding along the reaction coordinate (RC). The PMF highlights key thermodynamic properties of the binding process:

- **ΔG_on_**: The free energy of association calculated from the PMF minimum, representing the stability of the bound state.
- **ΔG_D_:** The dissociation energy barrier, indicating the energy required to disrupt the complex.
- **ΔG_off_:** The free energy difference between the bound and dissociated states. The binding events are categorized based on molecular recognition of PP1 (green-shaded region) and actin (red-shaded area), demonstrating the distinct energy landscapes associated with these binding interactions. **(b)** Quantitative parameters derived from the PMF profile for eIF2α binding. The free energy of association (**ΔG_on_**) and dissociation barrier (**ΔG_D_**) are used to infer the dissociation constant (**K_D_**) and approximate association (**k_on_**) and dissociation (**k_off_**) rates, linking the molecular dynamics data to mesoscale assembly kinetics. **(c)** Structural recognition of PP1 and actin by GADD34. The free energy of binding (**ΔG** _eIF2α**:PP1**_) and dissociation constant (**K_D_**) for each interaction were derived from the same simulation, based on the analysis of hydrogen bonds and residue contacts involving eIF2α, PP1, and actin.The free energy of binding (**ΔG** _eIF2α**:PP1**_) and dissociation constant (**K_D_**) for each interaction are shown, indicating that PP1 recognition (**ΔG**_eIF2α **:PP1**_ **= -50.189 kJ/mol, K_D_ = 3.42nM**) is more substantial compared to actin recognition (**ΔG** _eIF2α**:actin**_ **= -47.3927 kJ/mol**, **K_D_ = 10.01 nM**). This analysis highlights the hierarchical nature of molecular recognition in the PP1-GADD34-eIF2α complex assembly, with distinct energy landscapes and interaction affinities for PP1 and actin binding.

### GADD34 Unfolding and PP1 Binding

To further investigate how the conformational dynamics of GADD34 affect PP1 holoenzyme stability, a dedicated Umbrella Sampling setup was used to recapitulate the unbinding process of PP1 from GADD34, as shown in **Figure 3**. The PMF profile for GADD34 unbinding (**Figure 3a**) shows a stepwise unfolding process with distinct energy barriers that align with specific structural transitions. The dissociation barrier (**ΔG_PP1_**) delineates the stability of the bound state, while the **PMF_unfold_** provides insights into the energetic costs associated with GADD34’s conformational flexibility. Structural characterization along the reaction coordinate (**Figure 3c**) identified three intermediate states during the unfolding process: (**I**) the native folded conformation, where **GADD34** retains most of its native contacts; (**II**) a partially unfolded intermediate, where key binding regions are exposed to facilitate interaction with **PP1**; and (**III**) the fully unfolded state, in which GADD34 adopts a fully extended conformation. The free energy of binding between GADD34 and PP1 (**ΔG_PP1_ = 40.661 kJ/mol, K_D_ =0.138 M**) further supports the hypothesis that the GADD34 PP1-binding motif undergoes conformational transitions that enhance its molecular recognition capabilities.

**Figure 3.**
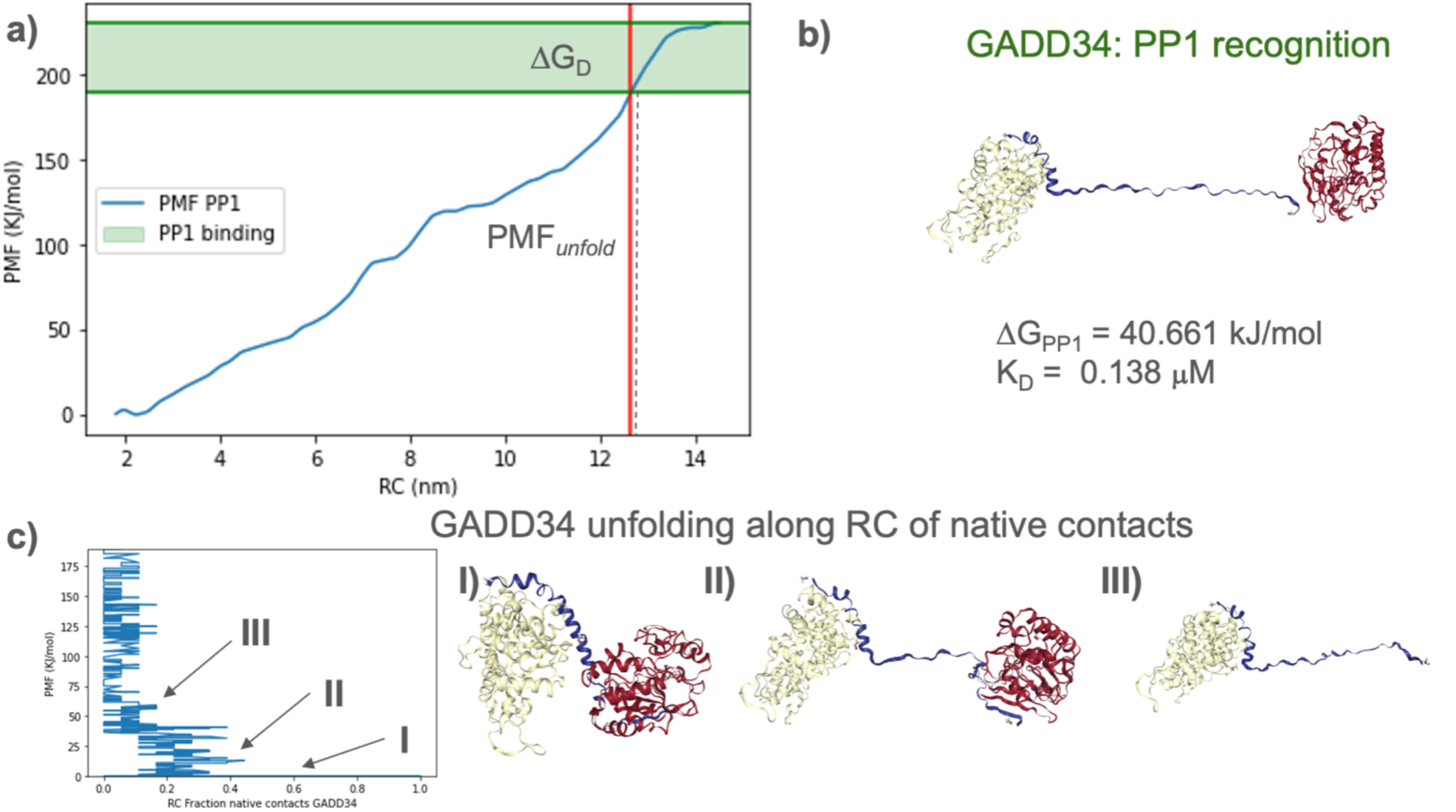
Energetic and structural analysis of GADD34 unfolding and its recognition of PP1. **(a)** Potential of Mean Force (PMF) profile for the interaction between GADD34 and PP1, calculated along the reaction coordinate (RC), defined as the distance between the center of masses of actin and PP1. The PMF highlights the unfolding process of GADD34 and its binding to PP1. The dissociation energy barrier (**ΔG_PP1_**) is marked, representing the free energy required to separate the complex. The green-shaded region corresponds to the stable binding state, while the ***PMF_unfold_*** indicates the unfolding energy of GADD34 when unbinds from PP1. **(b)** GADD34, PP1 recognition. The free energy of binding (**ΔG_PP1_** = 40.661 kJ/mol) and the dissociation constant (**K_D_** =0.138 *μ*M) are derived from the PMF profile, showing the moderate binding affinity between the two components. **(c)** The unfolding pathway of GADD34 along the RC is represented as a function of native contact fraction. Three key intermediates (I, II, III) are identified based on significant changes in PMF and structural features:

- **I:** Native folded state of GADD34, with most native contacts intact.
- **II:** Partially unfolded intermediate state with reduced native contacts, exposing binding regions for PP1.
- **III:** Fully unfolded state, characterized by minimal native contacts and extended conformation of GADD34.

This analysis provides a detailed view of GADD34 unfolding and binding to PP1, highlighting that the GADD34-PP1 binding domain is an intrinsically disordered protein (IDP).

### Bridging Energy Landscapes and Enzyme Kinetics: A PySB Model of GADD34-Mediated PP1 Regulation

To further elucidate how the folding states of GADD34 modulate the catalytic activity of the PP1 holoenzyme, we implemented an ordinary differential equation (ODE) model using PySB—a Python-based framework tailored for systems biology.^11^ The model was central to integrating and translating the detailed energy landscapes obtained from umbrella sampling into kinetic parameters that govern the dynamic assembly and activity of the PP1–GADD34–eIF2α complex. The computational advantage of this approach lies in its ability to incorporate energy constraints derived directly from umbrella sampling simulations by embedding the free energy barriers and intermediate state energetics into the ODE framework, yielding physically realistic rate constants and transition probabilities.^27,28^

### PySB Modeling and Fitness Landscape

As shown in **Figure 4**, the model accounts for the conformational dynamics of GADD34 by introducing an ad hoc variable, **ϕ**, which quantifies its folding state. Here, **ϕ** varies continuously from 0 to 1, corresponding to a fully unfolded to a fully native folded conformation, according to conformations observed in **Figure 3**. This variable is directly linked to the percentage of native contacts, defined as pairs of residues whose centers of mass remain within a threshold distance as in the native structure, indicating preserved tertiary interactions. It serves as a modulator in the kinetic equations, reflecting the allosteric influence of GADD34’s folding on PP1’s catalytic efficiency. As **ϕ** increases, indicative of a more ordered state either achieved through direct PP1 binding or actin-induced stabilization, the reorganization of PP1’s active site is optimized, thereby facilitating a more efficient dephosphorylation of eIF2α. Conversely, lower values of **ϕ** correspond to suboptimal catalytic alignment, which can reduce the reaction turnover. As shown in **Figure S1**, PP1 enzymatic activity is heavily affected by folding stability; indeed, the fitness landscape indicates that catalytic efficiency is maximized when Δ*G_fold_* is equal to 20 kJ/mol and **ϕ is** equal to 0.9. Consistently, the catalytic subunit PP1 concentration affects eIF2α enzymatic activity, as shown in **Figure S2**. Catalytic efficiency is reached for PP1 concentration equal to 800 mM and **ϕ** equal to 0.9.

**Figure 4.**
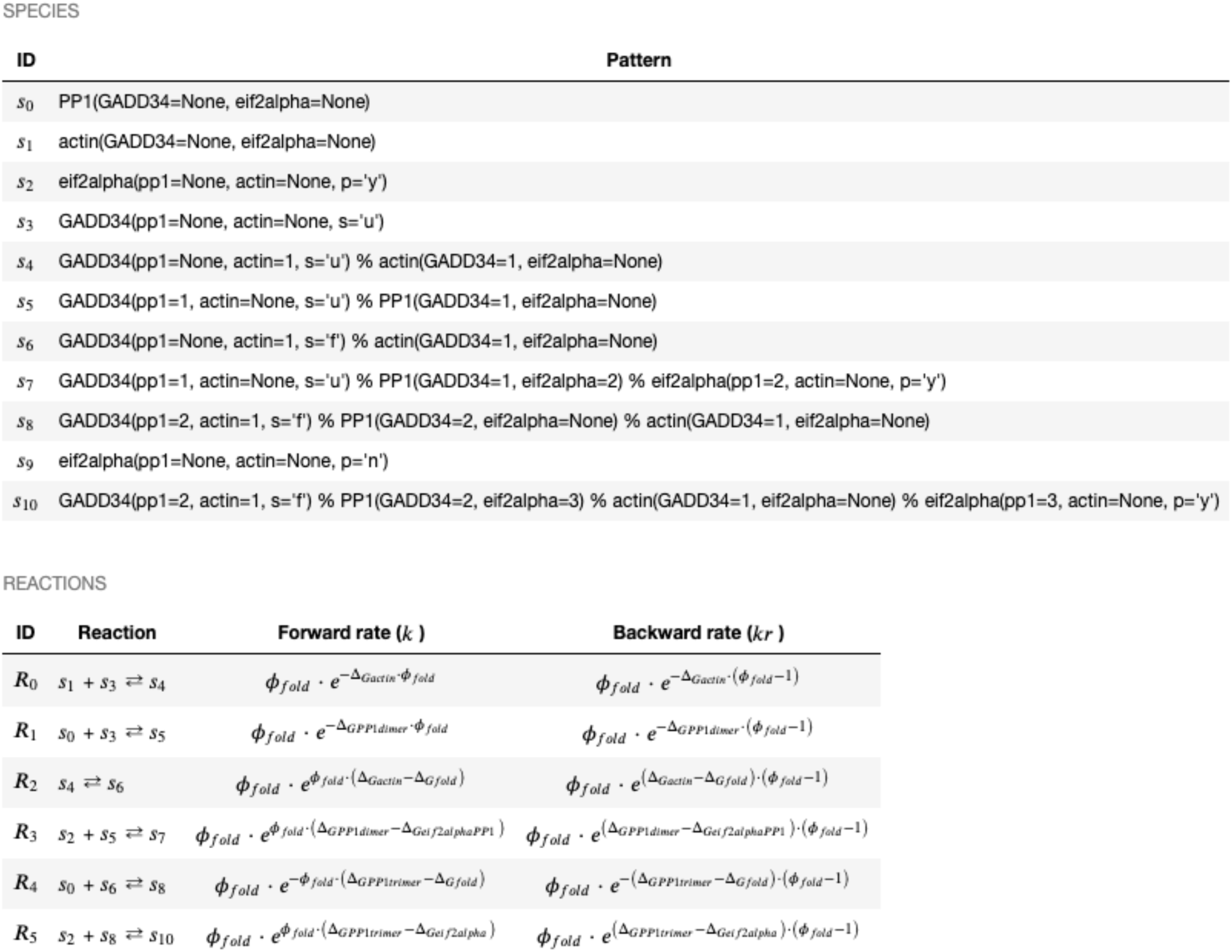
Table of context-dependent forward and reverse reaction rates. The reaction network for the PP1–GADD34–eIF2α–actin system was initially defined in PySB and subsequently translated into BioNetGen format. The top table lists all modeled species and their binding states, while the bottom table details the reaction definitions alongside the corresponding forward (k) and backward (kᵣ) rate constants. Notably, the parameter **ϕ**_fold_ accounts for the conformational dynamics of GADD34, thereby influencing the assembly and regulatory mechanisms of the PP1 holoenzyme.

Given that actin plays a key role in stabilizing the holoenzyme complex by binding to GADD34 and that GADD34’s binding affinity for PP1 is crucial for forming this complex, several simulations have been performed to identify the optimal combination of these two parameters. As shown in **Figure S3**, in these simulations, the model systematically varies both the actin–GADD34 and GADD34–PP1 binding affinities to determine which parameter sets yield the highest catalytic efficiency. The results underscore how the synergy between actin-induced stabilization and GADD34–PP1 interactions can significantly enhance the enzymatic output of the holoenzyme. Specifically, the maximum fitness is reached for Δ*G_Actin_* a Δ*G_PP_*_1_PP1 concentration equal to 3.3 kJ/mol and 38.8 kJ/mol, respectively. **Figure 5** further illustrates these effects by depicting the fitness landscape of eIF2α enzymatic activity at ϕ = 0.9. In panel (a), each point in the 3D surface represents a unique combination of GADD34’s binding affinity for actin (x-axis), binding affinity for PP1 (y-axis), and folding stability (z-axis or color scale), with the height/color indicating the predicted enzymatic activity (fitness). Panels (b) and (c) focus on cross-sections of the same parameter space—again at ϕ = 0.9—showing how varying binding affinities and folding stability influence the overall catalytic efficiency. Peaks in these surfaces correspond to parameter sets that maximize eIF2α dephosphorylation. This underscores how higher ϕ values (i.e., more ordered GADD34 states) can better align the active site residues in PP1, thereby promoting efficient substrate turnover.

**Figure 5.**
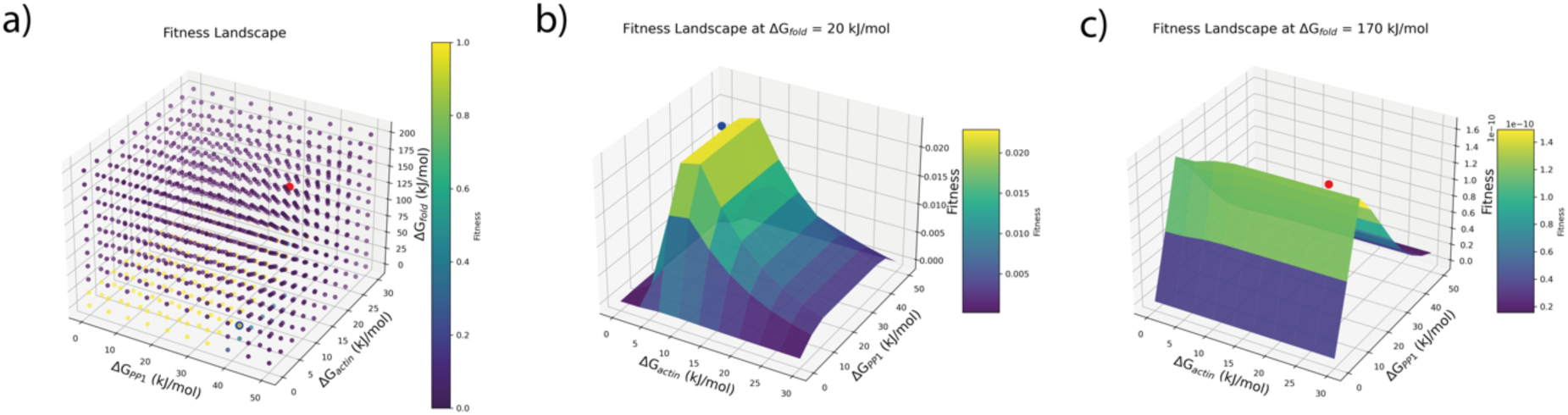
a) Three-dimensional plot illustrating the efficiency of PP1 complex enzymatic activity (color scale) as a function of Δ*G_PP_*_1_(x-axis), Δ*G_Actin_* (y-axis), and Δ*G_fold_* (z-axis). The maximum fitness is reached for Δ*G_Actin_*, Δ*G_PP_*_1_, Δ*G_fold_*, equal to 3.3 kJ/mol, 38.8 kJ/mol, and 20 kJ/mol, respectively. The red dot points to the conditions sampled through umbrella sampling MD simulations. b) Fitness landscape at Δ*G_fold_* equal to 20 kJ/mol. The blue dot represents the maximum fitness. c) Fitness landscape at Δ*G_fold_* equal to 170 kJ/mol. The red dot represents the maximum fitness.

### Synergistic Regulation of PP1 Holoenzyme Catalysis by Actin-Induced GADD34 Stabilization and PP1 Binding Dynamics

Ten simulations were performed under two energetic regimes to assess the impact of GADD34 folding stability and its binding affinities for PP1 and actin on the PP1 holoenzyme’s catalytic efficiency. The first regime employed constraints derived from Umbrella MD simulations (ΔG₍a₎ = 15.7 kJ/mol, ΔG₍PP1₎ = 40.661 kJ/mol, and ΔG₍fold₎ = 170 kJ/mol), while the second regime used optimized parameters based on maximum fitness (ΔG₍actin₎ = 3.3 kJ/mol, ΔG₍PP1₎ = 38.8 kJ/mol, and ΔG₍fold₎ = 20 kJ/mol). Both simulations were complemented by ODE simulations that systematically varied GADD34 folding states (0 < ϕ < 1.0).

**Figures S4** (non-optimized) and **S5** (optimized) illustrate how actin-induced stabilization and GADD34–PP1 binding collectively dictate catalytic output through distinct quantitative measures. In both regimes, optimal eIF2α dephosphorylation is observed under balanced interaction conditions, as indicated by the steady-state fraction of dephosphorylated eIF2α and the turnover rate assessed via the steady-state concentration of PP1–GADD34–eIF2α ternary complexes.

In the non-optimized regime (**Figure S4**), simulations indicate that folded GADD34 (ϕ > 0.5) is rapidly sequestered by actin, promoting the formation of an actin:GADD34 dimer that subsequently binds PP1 to generate the active PP1 holoenzyme complex responsible for efficient p-eIF2α dephosphorylation. In contrast, when GADD34 is predominantly unfolded (ϕ < 0.5), compensation occurs through the formation of GADD34:PP1 dimers—reflected by a higher channel capacity (0.0023 bit versus 0.0012 bit for the folded state, see **Figure 6a**)—and benefits from reduced activation energy for assembling the PP1–GADD34–P-eIF2α complex (as shown in **Figure 4),** resulting in faster dephosphorylation.

**Figure 6.**
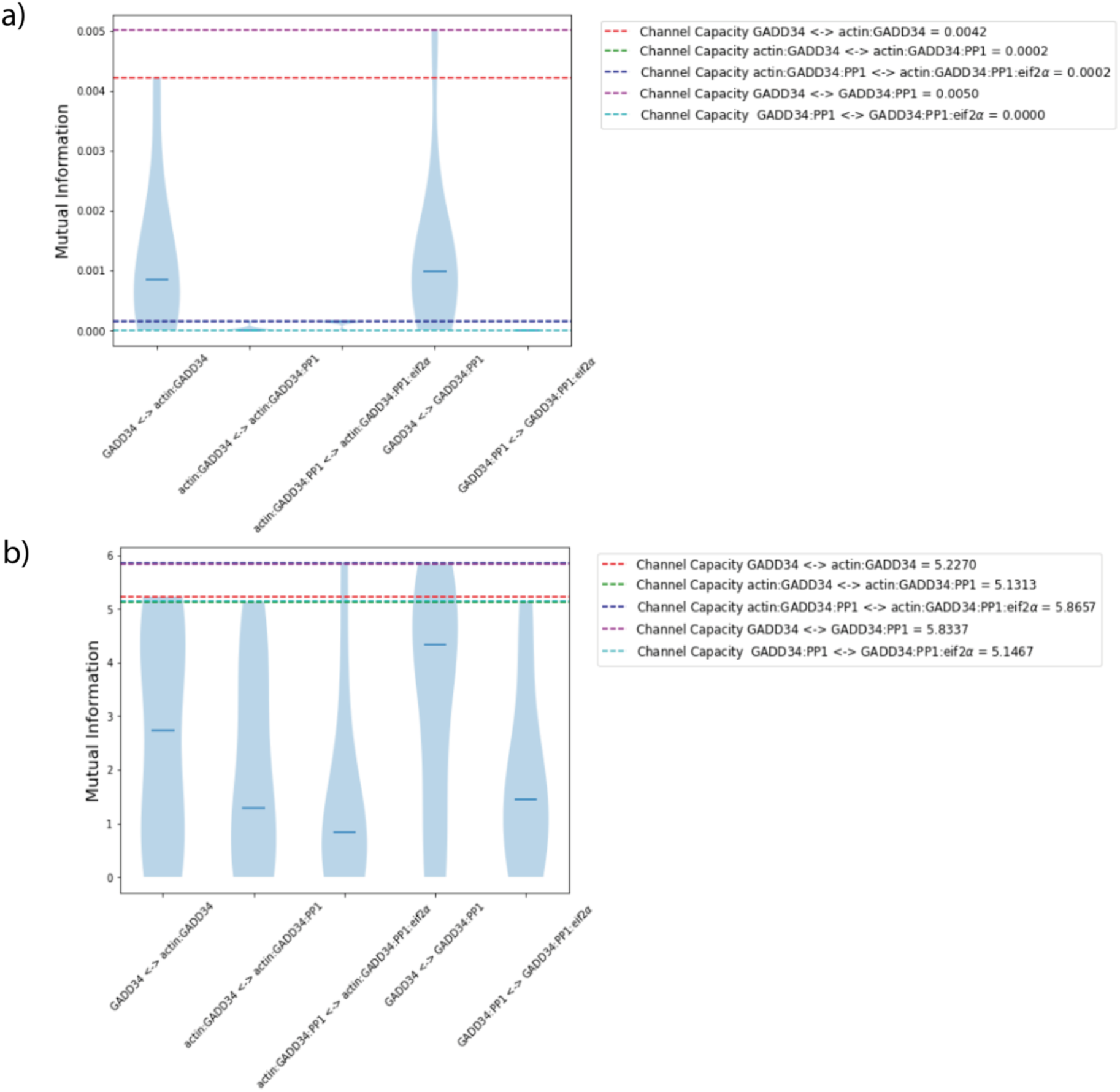
Comparison of Mutual Information and Channel Capacity in GADD34-Related Reactions Under Two Conditions. Panels **(a)** and **(b)** show violin plots of the mutual information (in bits) for GADD34-containing reaction pairs under two distinct conditions φ_fold was varied systematically and sampled discretely within a biologically relevant range to capture the system’s dynamic response without reliance on a fixed reference steady state. Mutual information between molecular species was estimated on frequency counts of min–max normalized time series. The dashed horizontal lines represent each reaction pair’s corresponding channel capacities (in bits). In panel **(a),** channel capacities range from approximately 0.0002 to 0.0021 bits, whereas in panel **(b),** they range from about 3.1467 to 5.6287 bits. In **a)**, the mutual information distributions lie well below the channel capacity lines (0.0002–0.0021 bits), indicating that, under these conditions, GADD34 folding and its interactions with actin and eIF2α do not utilize the full information-carrying potential. By contrast, **b)** reveals higher channel capacities (3.1467–5.6287 bits), and the corresponding mutual information distributions also increase, suggesting increased input–output sensitivity and greater capacity for control through GADD34 folding. These results underscore how changes in GADD34 folding and complex formation can significantly influence the system’s information flow efficiency.

In the optimized regime **(Figure S5**), both folded and unfolded GADD34 are efficiently sequestered by actin, facilitating the formation of actin:GADD34 dimers that subsequently bind PP1. Higher channel capacity values (5.227 bits for the holoenzyme complex and 5.833 bits for the compensatory GADD34:PP1 dimer in the unfolded state, see **Figure 6b**) indicate enhanced system responsiveness and assembly efficiency.

These findings underscore that the highest catalytic output is achieved through a synergistic interplay between actin-induced stabilization and optimal GADD34–PP1 binding, with optimized energetic parameters markedly enhancing eIF2α dephosphorylation efficiency.

## Discussion

The results presented here offer a mechanistic view of how conformational regulation influences the assembly and catalytic activity of the PP1–GADD34–eIF2α complex. Central to these findings is that the conformational state of GADD34 modulates its binding interactions with PP1 and actin, thereby influencing holoenzyme catalytic efficiency during cellular stress. The study reveals that GADD34 samples conformational states that modulate its binding preferences. GADD34 requires a binding partner—PP1 or actin—in its disordered state to induce a more ordered conformation. When GADD34 binds PP1 via its intrinsically disordered RVxF motif, induced folding may enhance the spatial arrangement of PP1’s catalytic site, supporting recruitment and positioning of eIF2α for dephosphorylation. Alternatively, actin binding stabilizes GADD34’s folded state and reduces the energetic cost of folding, indirectly promoting PP1 binding via an allosteric mechanism.

Umbrella sampling simulations provided quantitative estimates of the thermodynamic landscape governing complex assembly. The strong thermodynamic driving force for eIF2α binding (ΔG = –89.53 kJ/mol) and modest dissociation barrier (ΔG‡ = 10.58 kJ/mol) suggest a complex that is thermodynamically stable yet dynamically reversible. The calculated KD from the PMF (∼0.67 fM) may represent an upper limit, as the definition of the reaction coordinate and sampling boundaries could overestimate binding stability. The energy landscapes characterized by the potential of mean force (PMF) profiles highlight discrete intermediate states in GADD34 unfolding, suggesting that subtle conformational transitions can significantly affect binding energetics. These results support the idea that partial folding or structural transitions in GADD34 modulate PP1’s catalytic output by altering the orientation and spatial alignment of catalytic residues.

Integrating umbrella sampling data with a PySB-based ODE model represents a novel approach to connecting microscopic energy landscapes with macroscopic enzyme kinetics. By introducing the continuous folding variable ϕ, the model effectively translates structural changes into quantitative predictions of catalytic activity. The simulations reveal that maximal dephosphorylation of eIF2α occurs when GADD34 is in a highly ordered state (ϕ ≈ 0.9) and when PP1 concentration is optimized, highlighting the sensitivity of the holoenzyme’s function to both binding affinity and folding stability.

The phase space analysis further indicates that the synergy between actin-induced stabilization and optimal GADD34–PP1 interactions are paramount for achieving high catalytic output. Comparative simulations under constrained and optimized energetic regimes highlight how alternative assembly routes can support catalytic activity under suboptimal folding conditions. Under non-optimized conditions, compensation occurs through forming alternative dimeric complexes, ensuring a baseline level of dephosphorylation despite suboptimal folding. In contrast, optimized energetic parameters increase channel capacity and enhance the system’s responsiveness, resulting in robust eIF2α dephosphorylation.

## Conclusions

In summary, this work elucidates a regulatory mechanism in which the conformational dynamics of GADD34 modulate PP1 holoenzyme assembly and function. The intrinsic disorder of GADD34, and its folding upon binding to PP1 or stabilization by actin, critically influence the catalytic efficiency of the complex. Umbrella sampling simulations show that molecular recognition is thermodynamically favorable and involves conformational intermediates, reflecting the dynamic nature of GADD34 folding. By integrating structural free energy landscapes with a PySB-based kinetic model, we connect atomistic transitions to enzymatic activity and define quantitative regimes where folding state and binding energetics jointly optimize function. The highest catalytic output emerges from the combined effects of actin-induced stabilization and favorable GADD34–PP1 interactions, indicating that precise energetic tuning enables robust eIF2α dephosphorylation. Together, these results demonstrate how conformational control governs complex assembly and activity and provide a generalizable modeling framework for disordered proteins in signaling networks.

## ASSOCIATED CONTENT

## Supporting Information

The **Supporting Information** files are available free of charge.

Two-dimensional heatmap illustrating the efficiency of PP1 complex enzymatic activity (color scale) as a function of Δ*G_fold_* and **ϕ** of GADD34 (Figure S1); Two-dimensional heatmap illustrating the efficiency of PP1 complex enzymatic activity (color scale) as a function of PP1 concentration and **ϕ** of GADD34 (Figure S2); Two-dimensional heatmap illustrating the efficiency of PP1 complex enzymatic activity, PP1 concentration equal to 800 mM, **ϕ** equal to 0.9, Δ*G_fold_* equal to 20 kJ/mol, (color scale) as a function of Δ*G_actin_* (x-axis) and Δ*G_PP_*_1_ (y-axis), relative to the binding of GADD34 to actin and PP1. (Figure S3); The time course of intermediate dimers that concur to the formation of PP1 holoenzyme and dephosphorylation of phosphorylated-eif2*α*, as a function of **ϕ,** given Δ*G_actin_* equally to 15.7 kJ/mol, Δ*G_PP_*_1_ equal to 40.661 kJ/mol, and Δ*G_fold_* equal to 170 kJ/mol (Figure S4); The time course of intermediate dimers that concur to the formation of PP1 holoenzyme and dephosphorylation of phosphor-eif2*α*, as a function of **ϕ,** given Δ*G_actin_* equally to 3.3 kJ/mol, Δ*G_PP_*_1_equal to 38.8 kJ/mol, and Δ*G_fold_* equal to 20 kJ/mol. (Figure S5); Parameter used in the PySB model for the PP1–GADD34–eIF2α–actin system. Two values are shown: the “Energy Constrained State” and the “Optimal State,”(Table S1).

## Data and Software Availability

The PySB script, model details, and parameters used for the rule-based kinetic simulations are provided as Supporting Information. The molecular dynamics (MD) and umbrella sampling simulations were conducted using the open-source software GROMACS. Step-by-step simulation protocols required to reproduce the MD trajectories and potential of mean force (PMF) calculations are detailed in the Materials and Methods section. Mutual information analyses were performed using the PyInform library.

## AUTHOR INFORMATION

## Author Contributions

C.F.L. supervised the research. F.F., Z.A.L., and C.F.L. conceived the ideas. F.F. developed the methods and performed the simulations and computations. B.B. performed the simulations and computations. M.I. and M.O. helped generate the ODE-based models. The manuscript was written through contributions of all authors.

## ACKNOWLEDGMENT

The authors would like to thank Mauro Costa Mattioli and Lucas Reineke for their insightful conversations and critical feedback on this work and integrated stress response modeling.

## ABBREVIATIONS

GADD34: Growth Arrest and DNA-Damage-inducible protein 34
PP1: Protein Phosphatase-1
eIF2α: eukaryotic initiation factor 2 subunit alpha
MD: molecular dynamics.

---

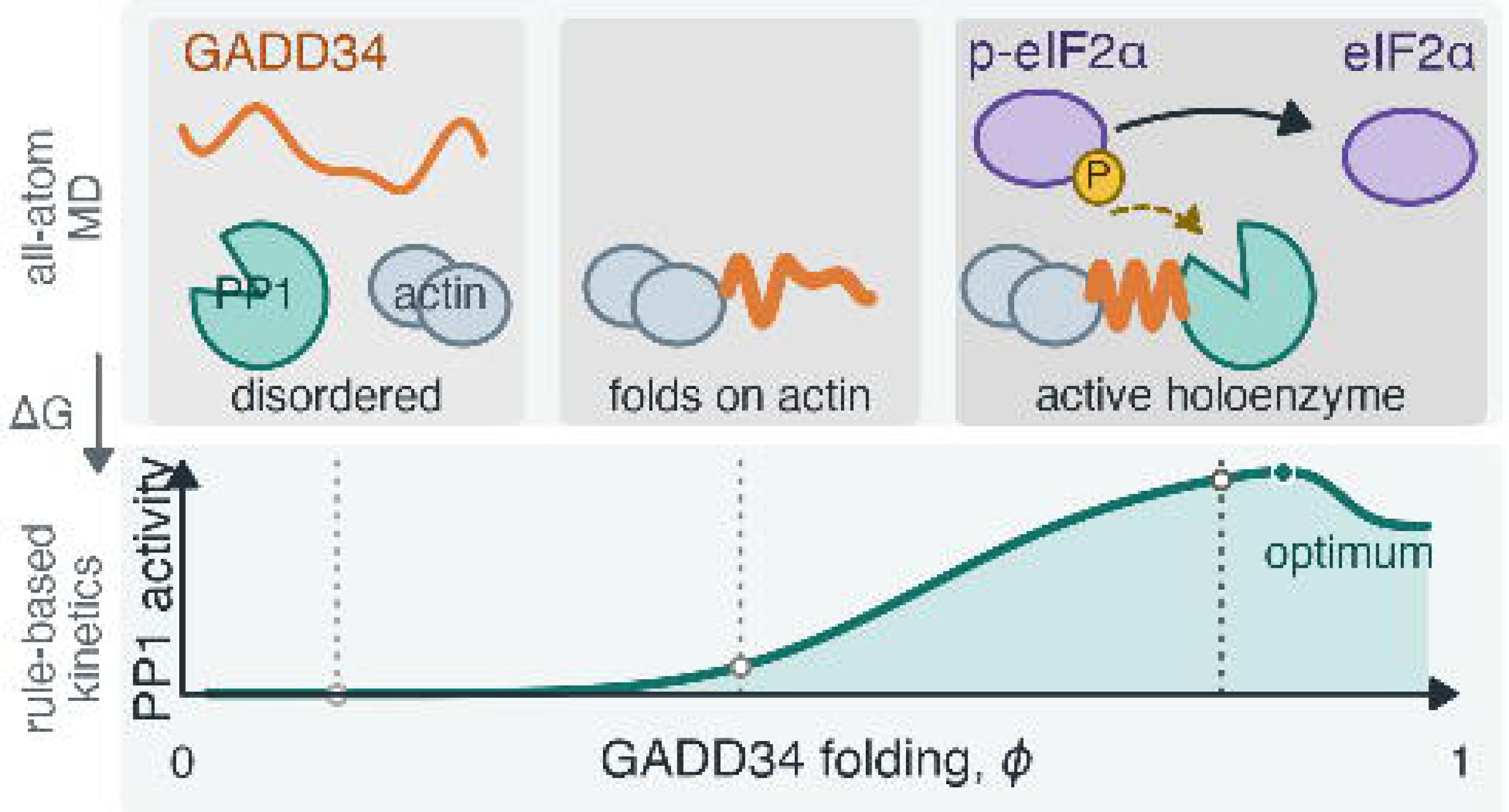

## REFERENCES

(1) Wang, Y. P.; Lei, Q. Y. Metabolite Sensing and Signaling in Cell Metabolism. Signal Transduction and Targeted Therapy 2018, 3 (1), 1–9. 10.1038/s41392-018-0024-7.

(2) Hibino, K.; Shibata, T.; Yanagida, T.; Sako, Y. Activation Kinetics of RAF Protein in the Ternary Complex of RAF, RAS-GTP, and Kinase on the Plasma Membrane of Living Cells: Single-Molecule Imaging Analysis. Journal of Biological Chemistry 2011, 286 (42), 36460–36468. 10.1074/jbc.M111.262675.

(3) Stewart, C. R.; Stuart, L. M.; Wilkinson, K.; Van Gils, J. M.; Deng, J.; Halle, A.; Rayner, K. J.; Boyer, L.; Zhong, R.; Frazier, W. A.; Lacy-Hulbert, A.; Khoury, J. El; Golenbock, D. T.; Moore, K. J. CD36 Ligands Promote Sterile Inflammation through Assembly of a Toll-like Receptor 4 and 6 Heterodimer. Nature Immunology 2010, 11 (2), 155–161. 10.1038/ni.1836.

(4) Pak, A. J.; Voth, G. A. Advances in Coarse-Grained Modeling of Macromolecular Complexes. Current Opinion in Structural Biology. Elsevier Ltd October 1, 2018, pp 119–126. 10.1016/j.sbi.2018.11.005.

(5) Freeman, R.; Boekhoven, J.; Dickerson, M. B.; Naik, R. R.; Stupp, S. I. Biopolymers and Supramolecular Polymers as Biomaterials for Biomedical Applications. MRS Bulletin 2015, 40 (12), 1089–1100. 10.1557/mrs.2015.270.

(6) van Vliet, A. R.; Giordano, F.; Gerlo, S.; Segura, I.; Van Eygen, S.; Molenberghs, G.; Rocha, S.; Houcine, A.; Derua, R.; Verfaillie, T.; Vangindertael, J.; De Keersmaecker, H.; Waelkens, E.; Tavernier, J.; Hofkens, J.; Annaert, W.; Carmeliet, P.; Samali, A.; Mizuno, H.; Agostinis, P. The ER Stress Sensor PERK Coordinates ER-Plasma Membrane Contact Site Formation through Interaction with Filamin-A and F-Actin Remodeling. Molecular Cell 2017, 65 (5), 885–899.e6. 10.1016/j.molcel.2017.01.020.

(7) Kmiecik, S.; Kolinski, A. Simulation of Chaperonin Effect on Protein Folding: A Shift from Nucleation - Condensation to Framework Mechanism. Journal of the American Chemical Society 2011, 133 (26), 10283–10289. 10.1021/ja203275f.

(8) Shammas, S. L.; Crabtree, M. D.; Dahal, L.; Wicky, B. I. M.; Clarke, J. Insights into Coupled Folding and Binding Mechanisms from Kinetic Studies *. Journal of Biological Chemistry 2016, 291 (13), 6689–6695. 10.1074/jbc.R115.692715.

(9) Souza, P. C. T.; Thallmair, S.; Conflitti, P.; Ramírez-Palacios, C.; Alessandri, R.; Raniolo, S.; Limongelli, V.; Marrink, S. J. Protein–Ligand Binding with the Coarse-Grained Martini Model. Nature Communications 2020, 11 (1). 10.1038/s41467-020-17437-5.

(10) Qian, H.; Beard, D. A.; Liang, S. D. Stoichiometric Network Theory for Nonequilibrium Biochemical Systems. European Journal of Biochemistry 2003, 270 (3), 415–421. 10.1046/j.1432-1033.2003.03357.x.

(11) Harris, L. A.; Nobile, M. S.; Pino, J. C.; Lubbock, A. L. R.; Besozzi, D.; Mauri, G.; Cazzaniga, P.; Lopez, C. F. GPU-Powered Model Analysis with PySB/CupSODA. Bioinformatics 2017, 33 (21), 3492–3494. 10.1093/bioinformatics/btx420.

(12) Klein, P.; Kallenberger, S. M.; Roth, H.; Roth, K.; Ly-Hartig, T. B. N.; Magg, V.; Aleš, J.; Talemi, S. R.; Qiang, Y.; Wolf, S.; Oleksiuk, O.; Kurilov, R.; Di Ventura, B.; Bartenschlager, R.; Eils, R.; Rohr, K.; Hamprecht, F. A.; Höfer, T.; Fackler, O. T.; Stoecklin, G.; Ruggieri, A. Temporal Control of the Integrated Stress Response by a Stochastic Molecular Switch. Science Advances 2022, 8 (12), 1–21. 10.1126/sciadv.abk2022.

(13) Costa-Mattioli, M.; Walter, P. The Integrated Stress Response: From Mechanism to Disease. Science (New York, N.Y.) 2020, 368 (6489). 10.1126/science.aat5314.

(14) Chiricotto, M.; Tran, T. T.; Nguyen, P. H.; Melchionna, S.; Sterpone, F.; Derreumaux, P. Coarse-Grained and All-Atom Simulations towards the Early and Late Steps of Amyloid Fibril Formation. Israel Journal of Chemistry 2017, 57 (7–8), 564–573. 10.1002/ijch.201600048.

(15) Wang, Z.; Huang, W.; Liu, M.; Kennel, S. J.; Wall, J. S.; Cheng, X. Computational Investigation of the Binding of a Designed Peptide to λ Light Chain Amyloid Fibril. Physical Chemistry Chemical Physics 2021, 23 (36), 20634–20644. 10.1039/d1cp01825f.

(16) Govind Kumar, V.; Polasa, A.; Agrawal, S.; Kumar, T. K. S.; Moradi, M. Binding Affinity Estimation from Restrained Umbrella Sampling Simulations. Nature Computational Science 2023, 3 (1), 59–70. 10.1038/s43588-022-00389-9.

(17) Yan, Y., Harding, H.P. & Ron, D. Higher-order phosphatase–substrate contacts terminate the integrated stress response. Nat Struct Mol Biol 2021, 28, 835–846

(18) Sankar, K.; Trainor, K.; Blazer, L. L.; Adams, J. J.; Sidhu, S. S.; Day, T.; Meiering, E.; Maier, J. K. X. A Descriptor Set for Quantitative Structure-Property Relationship Prediction in Biologics. Molecular Informatics 2022, 41 (9), 1–14. 10.1002/minf.202100240.

(19) Sankar, K.; Krystek, S. R.; Carl, S. M.; Day, T.; Maier, J. K. X. AggScore: Prediction of Aggregation-Prone Regions in Proteins Based on the Distribution of Surface Patches. Proteins: Structure, Function and Bioinformatics 2018, 86 (11), 1147–1156. 10.1002/prot.25594.

(20) Croll, T. I. ISOLDE: A Physically Realistic Environment for Model Building into Low-Resolution Electron-Density Maps. Acta Crystallographica Section D: Structural Biology 2018, 74 (6), 519–530. 10.1107/S2059798318002425.

(21) Choy, M. S.; Srivastava, G.; Robinson, L. C.; Tatchell, K.; Page, R.; Peti, W. The SDS22:PP1:I3 Complex: SDS22 Binding to PP1 Loosens the Active Site Metal to Prime Metal Exchange. Journal of Biological Chemistry 2024, 300 (1), 105515. 10.1016/j.jbc.2023.105515.

(22) McWhirter, C.; Lund, E. A.; Tanifum, E. A.; Feng, G.; Sheikh, Q. I.; Hengge, A. C.; Williams, N. H. Mechanistic Study of Protein Phosphatase-1 (PP1), a Catalytically Promiscuous Enzyme. Journal of the American Chemical Society 2008, 130 (41), 13673–13682. 10.1021/ja803612z.

(23) Fedoryshchak, R. O.; Přechová, M.; Butler, A.; Lee, R.; O’reilly, N.; Flynn, H.; Snijders, A. P.; Eder, N.; Ultanir, S.; Mouilleron, S.; Treisman, R. Molecular Basis for Substrate Specificity of the Phactr1/PP1 Phosphatase Holoenzyme. eLife 2020, 9, 1–78. 10.7554/ELIFE.61509.

(24) Ithuralde, R. E.; Roitberg, A. E.; Turjanski, A. G. Structured and Unstructured Binding of an Intrinsically Disordered Protein as Revealed by Atomistic Simulations. Journal of the American Chemical Society 2016, 138 (28), 8742–8751. 10.1021/jacs.6b02016.

(25) Chen, R.; Rato, C.; Yan, Y.; Crespillo-Casado, A.; Clarke, H. J.; Harding, H. P.; Marciniak, S. J.; Read, R. J.; Ron, D. G-Actin Provides Substrate-Specificity to Eukaryotic Initiation Factor2α Holophosphatases. eLife 2015, 2015 (4), 1–28. 10.7554/eLife.04871.

(26) Oliveira, M. M.; Mohamed, M.; Elder, M. K.; Banegas-Morales, K.; Mamcarz, M.; Lu, E. H.; Golhan, E. A. N.; Navrange, N.; Chatterjee, S.; Abel, T.; Klann, E. The Integrated Stress Response Effector GADD34 Is Repurposed by Neurons to Promote Stimulus-Induced Translation. Cell Reports 2024, 43 (2), 113670. 10.1016/j.celrep.2023.113670.

(27) Gerosa, L.; Chidley, C.; Fröhlich, F.; Sanchez, G.; Lim, S. K.; Muhlich, J.; Chen, J. Y.; Vallabhaneni, S.; Baker, G. J.; Schapiro, D.; Atanasova, M. I.; Chylek, L. A.; Shi, T.; Yi, L.; Nicora, C. D.; Claas, A.; Ng, T. S. C.; Kohler, R. H.; Lauffenburger, D. A.; Weissleder, R.; Miller, M. A.; Qian, W. J.; Wiley, H. S.; Sorger, P. K. Receptor-Driven ERK Pulses Reconfigure MAPK Signaling and Enable Persistence of Drug-Adapted BRAF-Mutant Melanoma Cells. Cell Systems 2020, 11 (5), 478–494.e9. 10.1016/j.cels.2020.10.002.

(28) Sekar, J. A. P.; Hogg, J. S.; Faeder, J. R. Energy-Based Modeling in BioNetGen. Proceedings - 2016 IEEE International Conference on Bioinformatics and Biomedicine, BIBM 2016 2017, 1460–1467. 10.1109/BIBM.2016.7822739.

