## Supplementary material for "Folding-Driven Control of the Functional PP1 Complex via Multiscale Modeling": Holoenzyme-formation-Execution-Modes-BioNetGen.ipynb

```
# %% [code]
from pysb import Model, Monomer, Parameter, Expression,  Rule, Observable, Initial, Annotation, EnergyPattern, ANY
from pysb.bng import generate_equations
from pysb.export import export
from pysb.core import as_complex_pattern, ComplexPattern
from sympy import exp, log, Symbol
import numpy as np
from pysb.macros import *

# %% [code]
Model()

# %% [code]
#The model consists of 4 different protein subunits (eif2alpha, PP1, actin, GADD34)
Monomer('eif2alpha', ['pp1', 'actin', 'p'], {'p': ['y', 'n']})
Monomer('PP1', ['GADD34', 'eif2alpha'])
Monomer('actin', ['GADD34', 'eif2alpha'])
Monomer('GADD34', ['pp1', 'actin', 's'], {'s': ['u', 'f']}) #GADD34 might exist in two alternative states folded and unfolded

# %% [code]
#Binding GADD34 to actin - this is the 1st reaction modeled in the network BINDING OF ACTIN TO GADD34 UNFOLDED
Parameter('GADD34_0',50)
Parameter('eif2alpha_0',2244) #Initial concentration of eif2alpha
Parameter('actin_0', 2800)
Parameter('PP1_0',800) #Initial concentration of PP1

Initial(PP1(GADD34=None,eif2alpha=None),PP1_0)
Initial(actin(GADD34=None,eif2alpha=None), actin_0)
Initial(eif2alpha(pp1=None,actin=None,p='y'), eif2alpha_0)
Initial(GADD34(pp1=None,actin=None,s='u'),GADD34_0)

Parameter('Delta_G_actin',15.70)
Parameter('phi_fold',1.0)

#convert kinetic parameters into energies for actin binding to GADD34
Expression('Gf_actin_GADD34', Delta_G_actin) #The energy Pattern is ln(KD)
Expression('Ea0_actin_GADD34', -log(phi_fold)) #The activation energy is strictly dependent on the folding distribution of GADD34
#define the energy pattern for the trimer
EnergyPattern('GADD34_bound_actin',actin(GADD34=1,eif2alpha=None)%GADD34(pp1=None,actin=1,s='u'), Gf_actin_GADD34)
Rule('GADD34_binds_actin',actin(GADD34=None,eif2alpha=None)+GADD34(pp1=None,actin=None,s='u') | 
     actin(GADD34=1,eif2alpha=None)%GADD34(pp1=None,actin=1,s='u'),phi_fold,Ea0_actin_GADD34,energy=True)

# %% [code]
Parameter('Delta_G_fold', 170.000)
Expression('Gf_fold', Delta_G_fold)
Expression('Ea0_fold', -log(phi_fold)) # The activation energy is strictly dependent on the folding distribution of GADD34
# define energy for GADD34 folding
EnergyPattern('GADD34_Folded', actin(GADD34=1, eif2alpha=None) % GADD34(pp1=None, actin=1, s='f'), Gf_fold)
#define rule for GADD34 folding
Rule('GADD34_folding', actin(GADD34=1, eif2alpha=None) % GADD34(pp1=None, actin=1, s='u') | actin(GADD34=1, eif2alpha=None) % GADD34(pp1=None, actin=1, s='f'), phi_fold, Ea0_fold, energy=True)

# %% [code]
#This is the 3rd reaction modeled in the network: BINDING OF GADD34:ACTIN to PP1
#Definition Rule with energy constraints given the gibbs free energy values obtained from MD simulations for PP1 binding GADD34
Delta_G_PP1_GADD34 = 40.661
Parameter('Delta_G_PP1_trimer', Delta_G_PP1_GADD34)

Expression('Gf_trimer', Delta_G_PP1_trimer) #unitless 
Expression('Ea0_trimer', - log(phi_fold)); #unitless 

#define the energy pattern for the trimer
EnergyPattern('PP1_trimer',actin(GADD34=1,eif2alpha=None)%GADD34(pp1=2,actin=1,s='f')%PP1(GADD34=2,eif2alpha=None),Gf_trimer)

#define PP1 binding to the dimer formed by GADD34 and actin
Rule('PP1_trimerization',actin(GADD34=1,eif2alpha=None)%GADD34(pp1=None,actin=1,s='f') + PP1(GADD34=None,eif2alpha=None)|
    actin(GADD34=1,eif2alpha=None)%GADD34(pp1=2,actin=1,s='f')%PP1(GADD34=2,eif2alpha=None),phi_fold,Ea0_trimer,energy=True)

# %% [code]
#convert kinetic parameters into energies for PP1 binding to GADD34
Parameter('Delta_G_PP1_dimer', Delta_G_PP1_GADD34)
Expression('Gf_dimer', Delta_G_PP1_dimer) 
Expression('Ea0_dimer', - log(phi_fold)) 
#define the energy pattern for the trimer
EnergyPattern('PP1_GADD34_dimer',GADD34(pp1=1,actin=None,s='u')%PP1(GADD34=1,eif2alpha=None),Gf_dimer)
#define PP1 binding to the dimer formed unfolded GADD34 
Rule('PP1_dimerization',GADD34(pp1=None,actin=None,s='u') + PP1(GADD34=None,eif2alpha=None)| GADD34(pp1=1,actin=None,s='u')%PP1(GADD34=1,eif2alpha=None),phi_fold,Ea0_dimer,energy=True)

# %% [code]
#This is the 2nd alternative reaction in modeled in the network: BINDING OF eif2alpha to unfolded GADDD34 bounds PP1
#Delta_G_eif2alpha_PP1 =  50189  #kJ/mol #dissociation energy eif2alpha from PP1
Parameter('Delta_G_eif2alpha_PP1', 50.189)
Expression('Gf_eif2alpha_PP1', Delta_G_eif2alpha_PP1)  
Expression('Ea0_eif2alpha_PP1', - log(phi_fold)); #unitless 
# define energy in bond between eif2alpha and actin
EnergyPattern('eif2alpha_PP1_binding',eif2alpha(pp1=2,actin=None,p='y')%GADD34(pp1=1,actin=None,s='u')%PP1(GADD34=1,eif2alpha=2),Gf_eif2alpha_PP1);
#define eif2alpha binding PP1 
Rule('eif2alpha_binds_PP1', eif2alpha(pp1=None,actin=None,p='y') + GADD34(pp1=1,actin=None,s='u')%PP1(GADD34=1,eif2alpha=None) | 
     eif2alpha(pp1=2,actin=None,p='y')%GADD34(pp1=1,actin=None,s='u')%PP1(GADD34=1,eif2alpha=2),phi_fold,Ea0_eif2alpha_PP1,energy=True)

# %% [code]
Parameter('Delta_G_eif2alpha',10.000)
Expression('Gf_Holoenzyme', Delta_G_eif2alpha) #unitless 
Expression('Ea0_Holoenzyme', - log(phi_fold)); #unitless 
#Definition Rule with energy constraints the values of gibbs free energy obtained from MD simulations for eif2alpha binding to PP1 trimer
EnergyPattern('Holoenzyme',actin(GADD34=1,eif2alpha=None)%GADD34(pp1=2,actin=1,s='f')%PP1(GADD34=2,eif2alpha=3)%eif2alpha(pp1=3,actin=None,p='y'),Gf_Holoenzyme);
#definition efi2alpha binding trimer
Rule('Holoenzyme_formation',actin(GADD34=1,eif2alpha=None)%GADD34(pp1=2,actin=1,s='f')%PP1(GADD34=2,eif2alpha=None) 
    + eif2alpha(pp1=None,actin=None,p='y') | actin(GADD34=1,eif2alpha=None)%GADD34(pp1=2,actin=1,s='f')%PP1(GADD34=2,eif2alpha=3)%eif2alpha(pp1=3,actin=None,p='y'),
    phi_fold,Ea0_Holoenzyme,energy=True) #Here we assume the distribution of folding is always equal to 1

# %% [code]
#Degradation anzd expression of GADD34 as derived from Ruggieri Article
Parameter('kdeg_GADD34',0.000309)##0.000309
Parameter('ksyn_GADD34',4.6e-5) ##4.6e-5
degrade(GADD34(pp1=None,actin=None,s='u'),kdeg_GADD34)
Rule('synth_GADD34u',eif2alpha(pp1=None,actin=None,p='y') >> GADD34(pp1=None,actin=None,s='u') + eif2alpha(pp1=None,actin=None,p='y'), ksyn_GADD34)

# %% [code]
# Define kinetic parameters and constants
Parameter('kcat_PP1', 2.23e6)   # intrinsic catalytic rate (s^-1 or appropriate units)
# Define the intrinsic catalytic energy (in kcal/mol)
Expression('Ecat0_catalysis', log(kcat_PP1))
# Define the effective catalytic energy that is modulated by the folding state (φ)
# ΔG_cat,eff = RT*ln(kcat_PP1) - RT*ln(phi_fold) = RT*ln(kcat_PP1/phi_fold)
Expression('Ecat_eff', Ecat0_catalysis - log(phi_fold))

# Define the energy pattern for the catalytic reaction.
# This assigns the effective catalytic energy to the reaction of dephosphorylating eif2alpha (p='n').
EnergyPattern('Enzyme_catalysis',
              eif2alpha(pp1=None, actin=None, p='n'),
              Ecat_eff)
Rule('Dephosphorylation_eif2alpha', actin(GADD34=1,eif2alpha=None)%GADD34(pp1=2,actin=1,s='f')%PP1(GADD34=2,eif2alpha=3)%eif2alpha(pp1=3,actin=None,p='y')>> 
    actin(GADD34=1,eif2alpha=None)%GADD34(pp1=2,actin=1,s='f')%PP1(GADD34=2,eif2alpha=None) + eif2alpha(pp1=None,actin=None,p='n'), Ecat_eff)
Rule('Dephosphorylation_eif2alpha_alt',eif2alpha(pp1=2,actin=None,p='y')%GADD34(pp1=1,actin=None,s='u')%PP1(GADD34=1,eif2alpha=2) >>
     eif2alpha(pp1=None,actin=None,p='n') + GADD34(pp1=1,actin=None,s='u')%PP1(GADD34=1,eif2alpha=None), Ecat_eff)

# %% [code]
Observable('ppp1r15a',GADD34(pp1=None,actin=None,s='u'))
Observable('GADD34_actin',actin(GADD34=1,eif2alpha=None)%GADD34(pp1=None,actin=1,s='u'))
Observable('GADD34_folded',actin(GADD34=1, eif2alpha=None) % GADD34(pp1=None, actin=1, s='f'))
Observable('PP1_trimer_folded',actin(GADD34=1,eif2alpha=None)%GADD34(pp1=2,actin=1,s='f')%PP1(GADD34=2,eif2alpha=None))
Observable('holoenzyme',actin(GADD34=1,eif2alpha=None)%GADD34(pp1=2,actin=1,s='f')%PP1(GADD34=2,eif2alpha=3)%eif2alpha(pp1=3,actin=None,p='y'))
Observable('GADD34_PP1', GADD34(pp1=1,actin=None,s='u')%PP1(GADD34=1,eif2alpha=None))
Observable('PP1_trimer_formed',eif2alpha(pp1=2,actin=None,p='y')%GADD34(pp1=1,actin=None,s='u')%PP1(GADD34=1,eif2alpha=2))

Observable('phospho_eif2alpha',eif2alpha(pp1=None,actin=None,p='y'))
Observable('dephospho_eif2alpha',eif2alpha(pp1=None,actin=None,p='n'))

#Channel 1: GADD34 -> GADD34:actin -> PP1:GADD34:actin -> PP1:GADD34:actin:p-eIF2alpha
#Channel 2: GADD34 -> GADD34:PP1 -> GADD34:PP1:eif2alpha

# %% [code]
## Visualize equation and species of the model
model.parameters

# %% [code]
import pandas as pd
import matplotlib.pyplot as plt
# Extract parameters from the model
data = [(p.name, p.value) for p in model.parameters]
df = pd.DataFrame(data, columns=['Parameter', 'Value'])

# Set up the matplotlib figure
fig, ax = plt.subplots(figsize=(6, 3))
ax.axis('tight')
ax.axis('off')

# Create the table and center-align the text
table = ax.table(cellText=df.values,
                 colLabels=df.columns,
                 cellLoc='center',
                 loc='center')

# Improve table aesthetics
table.auto_set_font_size(False)
table.set_fontsize(12)
table.scale(1.2, 1.2)

# Bold the header cells (column names)
for key, cell in table.get_celld().items():
    # Header cells are in the first row (row index 0)
    if key[0] == 0:
        cell.get_text().set_fontweight('bold')

# Optionally, add a title
#plt.title("PySB Model Parameters", fontsize=14)

# Save the figure with bbox_inches='tight' to capture the full table
plt.savefig("pysb_parameters.png", bbox_inches='tight')
plt.show()

# %% [code]
from util_display import display_model_info

# generate the model equations
generate_equations(model)

#display model informations
display_model_info(model)

#save the generated model in PySB and BNG format
generated_model_code = export(model, 'pysb_flat')
with open(model.name+'.py', 'wt') as f:
    f.write(generated_model_code);

generated_model_code = export(model, 'bngl')
with open(model.name+'.bngl', 'wt') as f:
    f.write(generated_model_code);

# %% [code]
#Adjust size of the table
from util_display import format_species_reactions, display_table
import pandas as pd
# Prevent pandas from truncating long LaTeX expressions when rendering.
pd.options.display.max_colwidth=None

#obtain dataframe with math latex expression visualization for species and reactions
(speciesdisp, reactionsdisp)=format_species_reactions(model);
#display species with the right style
display_table(speciesdisp, caption='SPECIES')

#display reactions in the same order as Reaction List S3 in original publication 
#reaction_order=[0,1,2,4,5,5]
display_table(reactionsdisp, caption='REACTIONS')
#display_table(reactionsdisp.iloc[reaction_order], caption='REACTIONS')

# %% [markdown]
## Fitness landscapes PP1 holoenzyme activities

# %% [markdown]
#### Fitness landscapes co-depency phi-fold and Delta_G_fold

# %% [code]
import numpy as np
from pysb.simulator import ScipyOdeSimulator
import matplotlib.pyplot as plt

def update_initial_conditions(model, initial_conditions):
    for monomer, value in initial_conditions.items():
        model.parameters[f"{monomer}_0"].value = value

def update_folding_parameters(model, folding_conditions):
    """
    Update folding-related parameters.
    Expects a dictionary with keys matching parameter names in the model,
    for example: {'phi_fold': value, 'Delta_G_fold': value}
    """
    for param, value in folding_conditions.items():
        model.parameters[param].value = value

# Time span for simulation (e.g., 4 days in seconds)
tspan = np.linspace(0, 345600, 115200)

# Define ranges for phi_fold (folding state) and Delta_G_fold.
# Here, phi_fold varies from 0 to 1 (11 intervals),
# and Delta_G_fold varies over an example range (e.g., 150 to 200 kcal/mol) in 11 intervals.
phi_vals = np.linspace(0, 1, 11)          # φ: 0, 0.1, 0.2, ..., 1.0
deltaG_vals = np.linspace(0, 200, 11)     # ΔG_fold in kcal/mol (example range)

# Fixed initial conditions for species that are not varied:
initial_conditions = {
    'eif2alpha': 2244,  # constant value (adjust as needed)
    'actin': 2400,      # constant value
    'GADD34': 40,       # fixed GADD34 concentration
    'PP1': 800          # fixed PP1 concentration
}
update_initial_conditions(model, initial_conditions)

# Prepare an empty fitness grid.
# Rows correspond to phi_fold values and columns to Delta_G_fold values.
fitness_grid = np.zeros((len(phi_vals), len(deltaG_vals)))

# Loop over phi_fold and Delta_G_fold values.
for i, phi_val in enumerate(phi_vals):
    for j, deltaG_val in enumerate(deltaG_vals):
        # Update the folding parameters in the model.
        folding_conditions = {
            'phi_fold': phi_val,
            'Delta_G_fold': deltaG_val
        }
        update_folding_parameters(model, folding_conditions)
        
        # Create a new simulator instance (using LSODA with specified tolerances)
        simulator = ScipyOdeSimulator(
            model,
            tspan=tspan,
            integrator_options={'rtol': 1e-12, 'atol': 1e-12}
        )
        sim_result = simulator.run()
        
        # Extract observables at the final time point.
        # (Ensure that your model defines these observables.)
        # For example:
        #   Observable('phospho_eif2alpha', eif2alpha(pp1=None, actin=None, p='y'))
        #   Observable('dephospho_eif2alpha', eif2alpha(pp1=None, actin=None, p='n'))
        obs_y = sim_result.observables['phospho_eif2alpha'][0]
        obs_n = sim_result.observables['dephospho_eif2alpha'][-1]
        
        # Compute fitness as the ratio: (dephospho)/(phospho).
        fitness = obs_n / obs_y if obs_y != 0 else 0
        fitness_grid[i, j] = fitness

# Create a meshgrid for plotting: ΔG_fold on the x-axis and phi_fold on the y-axis.
DeltaG_mesh, phi_mesh = np.meshgrid(deltaG_vals, phi_vals)

# Plot the fitness landscape as a filled contour plot.
plt.figure(figsize=(8, 6))
contour = plt.contourf(DeltaG_mesh, phi_mesh, fitness_grid, levels=50, cmap='viridis')
plt.xlabel('Delta_G_fold (kcal/mol)')
plt.ylabel('phi_fold (Folding State of GADD34)')
plt.title('Fitness Landscape\n[dephospho_eif2alpha] / [phospho_eif2alpha]')
plt.colorbar(contour, label='Fitness')
plt.show()

# %% [code]
from numpy import *
fitness_grid
where_are_NaNs = isnan(fitness_grid)
fitness_grid[where_are_NaNs] = 0
plt.figure(figsize=(8, 6))
contour = plt.contourf(DeltaG_mesh, phi_mesh, fitness_grid, levels=50, cmap='viridis')
plt.xlabel('$\Delta G_{fold}$ (kJ/mol)',fontsize=16)
plt.xticks(fontsize=16)
plt.yticks(fontsize=16)
plt.ylabel('$\phi_{fold}$ (Folding State of GADD34)',fontsize=16)
plt.title('Fitness Landscape',fontsize=16)
plt.colorbar(contour, label='Fitness')
plt.savefig('Figure-5.png',dpi=300)
plt.show()

# %% [code]
# Find the index of the maximum value in the flattened array
flat_index = np.argmax(fitness_grid)

# Convert the flattened index to a 2D coordinate (row, column)
max_coord = np.unravel_index(flat_index, fitness_grid.shape)

print("The maximum value is at:", max_coord)

# %% [code]
phi_vals[9],deltaG_vals[1]

# %% [markdown]
### Fitness landscapes co-depency phi-fold and PP1, fixed Delta_G_fold¶

# %% [code]
model.parameters['Delta_G_fold'].value = 20
model.parameters['Delta_G_fold'].value

# %% [code]
import numpy as np
from pysb.simulator import ScipyOdeSimulator
import matplotlib.pyplot as plt

def update_initial_conditions(model, initial_conditions):
    for monomer, value in initial_conditions.items():
        model.parameters[f"{monomer}_0"].value = value

def update_kinetic_conditions(model, kinetic_conditions):
    for rate, value in kinetic_conditions.items():
        # This assumes the kinetic parameter is named "phi_<rate>" (e.g., phi_fold)
        model.parameters[f"phi_{rate}"].value = value

# Time span for simulation (e.g., 4 days in seconds)
tspan = np.linspace(0, 345600, 115200)

# Define ranges for phi_fold (folding state) and PP1 initial concentration.
phi_vals = np.linspace(0, 1, 11)      # 0, 0.1, 0.2, ..., 1.0 (folding fraction)
pp1_vals = np.linspace(50, 800, 10)     # PP1 concentration range

# Prepare an empty fitness grid. Rows correspond to phi_fold values, columns to PP1.
fitness_grid = np.zeros((len(phi_vals), len(pp1_vals)))

# Loop over phi_fold and PP1 values.
for i, phi_val in enumerate(phi_vals):
    for j, pp1_val in enumerate(pp1_vals):
        # Set constant initial conditions for other species:
        # e.g., eIF2α and actin remain constant, GADD34 is fixed.
        initial_conditions = {
            'eif2alpha': 2244,
            'actin': 2400,
            'GADD34': 40  # Fixed value for GADD34
        }
        # PP1 is varied.
        initial_conditions['PP1'] = pp1_val
        update_initial_conditions(model, initial_conditions)
        
        # Update the kinetic condition for the folding state.
        # This updates the parameter "phi_fold" in your model.
        kinetic_conditions = {'fold': phi_val}
        update_kinetic_conditions(model, kinetic_conditions)
        
        # Create a new simulator instance (using LSODA with specified tolerances)
        simulator = ScipyOdeSimulator(
            model,
            tspan=tspan,
            integrator_options={'rtol': 1e-12, 'atol': 1e-12}
        )
        sim_result = simulator.run()
        
        # Extract the observables at the final time point.
        # Ensure that your model defines these observables:
        #   Observable('dephos_eif2alpha_n', eif2alpha(pp1=None, actin=None, p='n'))
        #   Observable('dephos_eif2alpha_y', eif2alpha(pp1=None, actin=None, p='y'))
        obs_y = sim_result.observables['phospho_eif2alpha'][0]
        obs_n = sim_result.observables['dephospho_eif2alpha'][-1]
        
        # Compute fitness as the ratio. (Use np.nan if obs_y is zero.)
        fitness = obs_n / obs_y if obs_y != 0 else 0
        fitness_grid[i, j] = fitness

# Create a meshgrid for plotting: PP1 on the x-axis and phi_fold on the y-axis.
PP1_mesh, phi_mesh = np.meshgrid(pp1_vals, phi_vals)

# Plot the fitness landscape as a filled contour plot.
plt.figure(figsize=(8, 6))
contour = plt.contourf(PP1_mesh, phi_mesh, fitness_grid, levels=50, cmap='viridis')
plt.xlabel('PP1 Concentration')
plt.ylabel('phi_fold (Folding State of GADD34)')
plt.title('Fitness Landscape\n[dephos_eif2alpha (p="n")] / [dephos_eif2alpha (p="y")]')
plt.colorbar(contour, label='Fitness')
plt.show()

# %% [code]
from numpy import *
fitness_grid
where_are_NaNs = isnan(fitness_grid)
fitness_grid[where_are_NaNs] = 0
plt.figure(figsize=(8, 6))
contour = plt.contourf(PP1_mesh, phi_mesh, fitness_grid, levels=50, cmap='viridis')
plt.xlabel('PP1 Concentration',fontsize=16)
plt.xticks(fontsize=16)
plt.yticks(fontsize=16)
plt.ylabel('$\phi_{fold}$ (Folding State of GADD34)',fontsize=16)
plt.title('Fitness Landscape',fontsize=16)
plt.colorbar(contour, label='Fitness')
plt.savefig('Figure-S1.png',dpi=300)
plt.show()

# %% [code]
# Find the index of the maximum value in the flattened array
flat_index = np.argmax(fitness_grid)

# Convert the flattened index to a 2D coordinate (row, column)
max_coord = np.unravel_index(flat_index, fitness_grid.shape)

print("The maximum value is at:", max_coord)

# %% [code]
phi_vals[9],pp1_vals[9]

# %% [markdown]
#### Fitness landscapes co-depency Delta_G_actin and Delta_G_PP1, fixed Delta_G_fold, phi_fold and PP1¶

# %% [code]
model.parameters

# %% [code]
import numpy as np
from pysb.simulator import ScipyOdeSimulator
import matplotlib.pyplot as plt

def update_initial_conditions(model, initial_conditions):
    for monomer, value in initial_conditions.items():
        model.parameters[f"{monomer}_0"].value = value

def update_folding_parameters(model, folding_conditions):
    """
    Update folding-related parameters.
    Expects a dictionary with keys matching parameter names in the model,
    for example: {'phi_fold': value, 'Delta_G_fold': value}
    """
    for param, value in folding_conditions.items():
        model.parameters[param].value = value

# Time span for simulation (e.g., 4 days in seconds)
tspan = np.linspace(0, 345600, 115200)

# Define ranges for phi_fold (folding state) and Delta_G_fold.
# Here, phi_fold varies from 0 to 1 (11 intervals),
# and Delta_G_fold varies over an example range (e.g., 150 to 200 kcal/mol) in 11 intervals.
deltaG_PP1_vals = np.linspace(0, 50, 10)          # φ: 0, 0.1, 0.2, ..., 1.0
deltaG_actin_vals = np.linspace(0, 30, 10)     # ΔG_fold in kcal/mol (example range)

# Fixed initial conditions for species that are not varied:
initial_conditions = {
    'eif2alpha': 2244,  # constant value (adjust as needed)
    'actin': 2400,      # constant value
    'GADD34': 40,       # fixed GADD34 concentration
    'PP1': 800          # fixed PP1 concentration
}
update_initial_conditions(model, initial_conditions)

# Prepare an empty fitness grid.
# Rows correspond to phi_fold values and columns to Delta_G_fold values.
fitness_grid = np.zeros((len(deltaG_actin_vals), len(deltaG_PP1_vals)))

# Loop over phi_fold and Delta_G_fold values.
for i, deltaG_PP1_val in enumerate(deltaG_PP1_vals):
    for j, deltaG_actin_val in enumerate(deltaG_actin_vals):
        # Update the folding parameters in the model.
        folding_conditions = {
            'phi_fold': 0.9,
            'Delta_G_fold': 20,
            'Delta_G_actin': deltaG_actin_val,
            'Delta_G_PP1_trimer':deltaG_PP1_val,
            'Delta_G_PP1_dimer': deltaG_PP1_val
            
        }
        update_folding_parameters(model, folding_conditions)
        
        # Create a new simulator instance (using LSODA with specified tolerances)
        simulator = ScipyOdeSimulator(
            model,
            tspan=tspan,
            integrator_options={'rtol': 1e-12, 'atol': 1e-12}
        )
        sim_result = simulator.run()
        
        # Extract observables at the final time point.
        # (Ensure that your model defines these observables.)
        # For example:
        #   Observable('phospho_eif2alpha', eif2alpha(pp1=None, actin=None, p='y'))
        #   Observable('dephospho_eif2alpha', eif2alpha(pp1=None, actin=None, p='n'))
        obs_y = sim_result.observables['phospho_eif2alpha'][0]
        obs_n = sim_result.observables['dephospho_eif2alpha'][-1]
        
        # Compute fitness as the ratio: (dephospho)/(phospho).
        fitness = obs_n / obs_y if obs_y != 0 else 0
        fitness_grid[i, j] = fitness

# Create a meshgrid for plotting: ΔG_fold on the x-axis and phi_fold on the y-axis.
DeltaG_actin_mesh, DeltaG_PP1_mesh = np.meshgrid(deltaG_actin_vals, deltaG_PP1_vals)

# Plot the fitness landscape as a filled contour plot.
plt.figure(figsize=(8, 6))
contour = plt.contourf(DeltaG_actin_mesh, DeltaG_PP1_mesh, fitness_grid, levels=50, cmap='viridis')
plt.xlabel('Delta_G_actin (kJ/mol)')
plt.ylabel('Delta_G_PP1 (kJ/mol)')
plt.title('Fitness Landscape\n[dephospho_eif2alpha] / [phospho_eif2alpha]')
plt.colorbar(contour, label='Fitness')
plt.show()

# %% [code]
from numpy import *
# Create a meshgrid for plotting: ΔG_fold on the x-axis and phi_fold on the y-axis.
DeltaG_actin_mesh, DeltaG_PP1_mesh = np.meshgrid(deltaG_actin_vals, deltaG_PP1_vals)

# Plot the fitness landscape as a filled contour plot.
plt.figure(figsize=(8, 6))
where_are_NaNs = isnan(fitness_grid)
fitness_grid[where_are_NaNs] = 0
contour = plt.contourf(DeltaG_actin_mesh, DeltaG_PP1_mesh, fitness_grid, levels=50, cmap='viridis')
plt.xlabel('$\Delta G_{actin}$ (kJ/mol)',fontsize=16)
plt.ylabel('$\Delta G_{PP1}$ (kJ/mol)', fontsize=16)
# Add a red dot at a fixed position, e.g., at (0, 0)
plt.plot(15.7, 40.6, 'ro', markersize=8)  # 
plt.xticks(fontsize=16)
plt.yticks(fontsize=16)
plt.title('Fitness Landscape',fontsize=16)
plt.colorbar(contour, label='Fitness')
plt.savefig('Figure-S2.png',dpi=300)
plt.show()

# %% [code]


# %% [code]
#Find the index of the maximum value in the flattened array
flat_index = np.argmax(fitness_grid)

# Convert the flattened index to a 2D coordinate (row, column)
max_coord = np.unravel_index(flat_index, fitness_grid.shape)

print("The maximum value is at:", max_coord)

# %% [code]
deltaG_actin_vals[1],deltaG_PP1_vals[7]

# %% [markdown]
## Fitness landscapes co-depency Delta_G_actin and Delta_G_PP1, Delta_G_fold, fixed phi_fold = 0.9 and PP1 = 800 mM¶


# %% [code]
import numpy as np
from pysb.simulator import ScipyOdeSimulator
import matplotlib.pyplot as plt

def update_initial_conditions(model, initial_conditions):
    for monomer, value in initial_conditions.items():
        model.parameters[f"{monomer}_0"].value = value

def update_folding_parameters(model, folding_conditions):
    """
    Update folding-related parameters.
    Expects a dictionary with keys matching parameter names in the model,
    for example: {'phi_fold': value, 'Delta_G_fold': value}
    """
    for param, value in folding_conditions.items():
        model.parameters[param].value = value

# Time span for simulation (e.g., 4 days in seconds)
tspan = np.linspace(0, 345600, 115200)

# Define ranges for phi_fold (folding state) and Delta_G_fold.
# Here, phi_fold varies from 0 to 1 (11 intervals),
# and Delta_G_fold varies over an example range (e.g., 150 to 200 kcal/mol) in 11 intervals.
deltaG_PP1_vals = np.linspace(0, 50, 10)          # φ: 0, 0.1, 0.2, ..., 1.0
deltaG_actin_vals = np.linspace(0, 30, 10)     # ΔG_fold in kcal/mol (example range)
deltaG_vals = np.linspace(0, 200, 11) 
# Fixed initial conditions for species that are not varied:
initial_conditions = {
    'eif2alpha': 2244,  # constant value (adjust as needed)
    'actin': 2400,      # constant value
    'GADD34': 40,       # fixed GADD34 concentration
    'PP1': 800          # fixed PP1 concentration
}
update_initial_conditions(model, initial_conditions)

# Prepare an empty fitness grid.
# Rows correspond to phi_fold values and columns to Delta_G_fold values.
fitness_grid = np.zeros((len(deltaG_vals),len(deltaG_actin_vals), len(deltaG_PP1_vals)))

# Loop over phi_fold and Delta_G_fold values.
for i, delta_G_val in enumerate(deltaG_vals):
    for j, deltaG_actin_val in enumerate(deltaG_actin_vals):
        for k, deltaG_PP1_val in enumerate(deltaG_PP1_vals):
        # Update the folding parameters in the model.
            folding_conditions = {
                'phi_fold': 0.9,
                'Delta_G_fold': delta_G_val,
                'Delta_G_actin': deltaG_actin_val,
                'Delta_G_PP1_trimer':deltaG_PP1_val,
                'Delta_G_PP1_dimer': deltaG_PP1_val
            
            }
            update_folding_parameters(model, folding_conditions)
        
            # Create a new simulator instance (using LSODA with specified tolerances)
            simulator = ScipyOdeSimulator(
                model,
                tspan=tspan,
                integrator_options={'rtol': 1e-12, 'atol': 1e-12}
            )
            sim_result = simulator.run()
        
        # Extract observables at the final time point.
        # (Ensure that your model defines these observables.)
        # For example:
        #   Observable('phospho_eif2alpha', eif2alpha(pp1=None, actin=None, p='y'))
        #   Observable('dephospho_eif2alpha', eif2alpha(pp1=None, actin=None, p='n'))
            obs_y = sim_result.observables['phospho_eif2alpha'][0]
            obs_n = sim_result.observables['dephospho_eif2alpha'][-1]
        
            # Compute fitness as the ratio: (dephospho)/(phospho).
            fitness = obs_n / obs_y if obs_y != 0 else 0
            fitness_grid[i, j, k] = fitness


# %% [code]
import numpy as np
import matplotlib.pyplot as plt
from mpl_toolkits.mplot3d import Axes3D  # required for 3D plotting

# Assume these arrays and fitness_grid have been computed earlier in your code:
# deltaG_PP1_vals  (e.g., np.linspace(0, 50, 10))
# deltaG_actin_vals (e.g., np.linspace(0, 30, 10))
# deltaG_vals      (e.g., np.linspace(0, 200, 11))
# fitness_grid with shape (len(deltaG_vals), len(deltaG_actin_vals), len(deltaG_PP1_vals))
#   because your loops were:
#     for i, delta_G_val in enumerate(deltaG_vals):
#         for j, deltaG_actin_val in enumerate(deltaG_actin_vals):
#             for k, deltaG_PP1_val in enumerate(deltaG_PP1_vals):
#                 fitness_grid[i, j, k] = fitness

# Create a 3D meshgrid with the same axis order:
# First axis: deltaG_vals (ΔG_fold)
# Second axis: deltaG_actin_vals (ΔG_actin)
# Third axis: deltaG_PP1_vals (ΔG_PP1)
DeltaG_fold_mesh, DeltaG_actin_mesh, DeltaG_PP1_mesh = np.meshgrid(
    deltaG_vals, deltaG_actin_vals, deltaG_PP1_vals, indexing='ij'
)

# Replace NaNs in fitness_grid (if any) with zeros.
fitness_grid = np.nan_to_num(fitness_grid)

# Flatten the meshgrid arrays and the fitness grid for plotting.
# Note: x corresponds to ΔG_PP1, y to ΔG_actin, and z to ΔG_fold.
x = DeltaG_PP1_mesh.flatten()
y = DeltaG_actin_mesh.flatten()
z = DeltaG_fold_mesh.flatten()
c = fitness_grid.flatten()  # Fitness values for the colormap

# Create the 3D scatter plot.
fig = plt.figure(figsize=(10, 8))
ax = fig.add_subplot(111, projection='3d')
sc = ax.scatter(x, y, z, c=c, cmap='viridis', marker='o')

# Add a colorbar and set labels.
plt.colorbar(sc, ax=ax, label='Fitness')

# Plot highlighted points (using scatter for clarity)
ax.scatter([40.6], [15.7], [170], c='r', s=80, marker='o')  # Example red marker
ax.scatter([38.8], [3.3], [20], c='b', s=80, marker='o')     # Example blue marker

ax.set_xlabel('ΔG$_{PP1}$ (kJ/mol)', fontsize=16)
ax.set_ylabel('ΔG$_{actin}$ (kJ/mol)', fontsize=16)
ax.set_zlabel('ΔG$_{fold}$ (kJ/mol)', fontsize=16)
plt.title('Fitness Landscape', fontsize=16)
plt.savefig('Figure-5.png', dpi=300)
plt.show()


# %% [code]
#Find the index of the maximum value in the flattened array
flat_index = np.argmax(fitness_grid)

# Convert the flattened index to a 2D coordinate (row, column)
max_coord = np.unravel_index(flat_index, fitness_grid.shape)

print("The maximum value is at:", max_coord)

# %% [code]
deltaG_PP1_vals[1],deltaG_actin_vals[1],deltaG_vals[7]   # ΔG_fold in kcal/mol (example range)

# %% [code]
# From here run simulations with original set of parameters and set of parameters from sampling fitness landscape and varying phi-fold

# %% [code]
#Calculate only gains P-eif2alpha[0]/eif2alpha

# %% [code]
fitness_grid.shape

# %% [code]
deltaG_vals

# %% [code]
from mpl_toolkits.mplot3d import Axes3D  # Required for 3D plotting
import numpy as np
import matplotlib.pyplot as plt

# Assume these arrays and fitness_grid have been computed earlier:
# deltaG_PP1_vals  (e.g., np.linspace(0, 50, 10))
# deltaG_actin_vals (e.g., np.linspace(0, 30, 10))
# deltaG_vals      (e.g., np.linspace(0, 200, 11))
# fitness_grid with shape (len(deltaG_vals), len(deltaG_actin_vals), len(deltaG_PP1_vals))
# (with the ordering: [ΔG_fold, ΔG_actin, ΔG_PP1])

# Select a fixed ΔG_fold slice, e.g., the middle value:
k_index = 2
fitness_slice = fitness_grid[k_index, :, :]  # shape: (len(deltaG_actin_vals), len(deltaG_PP1_vals))

# Create a 2D meshgrid for ΔG_actin (x-axis) and ΔG_PP1 (y-axis).
# Using indexing='ij' ensures that the output arrays have shape
# (len(deltaG_actin_vals), len(deltaG_PP1_vals)) to match fitness_slice.
DeltaG_actin_mesh, DeltaG_PP1_mesh = np.meshgrid(deltaG_actin_vals, deltaG_PP1_vals, indexing='ij')

# Set up the 3D plot.
fig = plt.figure(figsize=(10, 8))
ax = fig.add_subplot(111, projection='3d')

# Plot the surface with:
#   x = ΔG_actin, y = ΔG_PP1, z = fitness value.
surf = ax.plot_surface(DeltaG_actin_mesh, DeltaG_PP1_mesh, fitness_slice,
                       cmap='viridis', edgecolor='none', alpha=0.9)

# Set axis labels and title.
ax.set_xlabel('ΔG$_{actin}$ (kJ/mol)', fontsize=16)
ax.set_ylabel('ΔG$_{PP1}$ (kJ/mol)', fontsize=16)
ax.set_zlabel('Fitness', fontsize=16)
ax.set_title(f'Fitness Landscape at ΔG$_{{fold}}$ = 20 kJ/mol', fontsize=16)

# Highlight a specific point on the surface (example coordinates).
ax.scatter(3.5, 38.8, 0.02, c='b', s=80, marker='o')

# Add a colorbar which maps values to colors.
fig.colorbar(surf, shrink=0.5, aspect=5, label='Fitness')
plt.savefig('Figure-5a.png', dpi=300)
plt.show()


# %% [code]
from mpl_toolkits.mplot3d import Axes3D  # Required for 3D plotting
import numpy as np
import matplotlib.pyplot as plt

# Assume these arrays and fitness_grid have been computed earlier:
# deltaG_PP1_vals  (e.g., np.linspace(0, 50, 10))
# deltaG_actin_vals (e.g., np.linspace(0, 30, 10))
# deltaG_vals      (e.g., np.linspace(0, 200, 11))
# fitness_grid with shape (len(deltaG_vals), len(deltaG_actin_vals), len(deltaG_PP1_vals))
# (with the ordering: [ΔG_fold, ΔG_actin, ΔG_PP1])

# Select a fixed ΔG_fold slice, e.g., the middle value:
k_index = 10
fitness_slice = fitness_grid[k_index, :, :]  # shape: (len(deltaG_actin_vals), len(deltaG_PP1_vals))

# Create a 2D meshgrid for ΔG_actin (x-axis) and ΔG_PP1 (y-axis).
# Using indexing='ij' ensures that the output arrays have shape
# (len(deltaG_actin_vals), len(deltaG_PP1_vals)) to match fitness_slice.
DeltaG_actin_mesh, DeltaG_PP1_mesh = np.meshgrid(deltaG_actin_vals, deltaG_PP1_vals, indexing='ij')

# Set up the 3D plot.
fig = plt.figure(figsize=(10, 8))
ax = fig.add_subplot(111, projection='3d')

# Plot the surface with:
#   x = ΔG_actin, y = ΔG_PP1, z = fitness value.
surf = ax.plot_surface(DeltaG_actin_mesh, DeltaG_PP1_mesh, fitness_slice,
                       cmap='viridis', edgecolor='none', alpha=0.9)

# Set axis labels and title.
ax.set_xlabel('ΔG$_{actin}$ (kJ/mol)', fontsize=16)
ax.set_ylabel('ΔG$_{PP1}$ (kJ/mol)', fontsize=16)
ax.set_zlabel('Fitness', fontsize=16)
ax.set_title(f'Fitness Landscape at ΔG$_{{fold}}$ = 170 kJ/mol', fontsize=16)

# Highlight a specific point on the surface (example coordinates).
ax.scatter(15.7, 39.6, 0.9e-10, c='r', s=80, marker='o')

# Add a colorbar which maps values to colors.
fig.colorbar(surf, shrink=0.5, aspect=5, label='Fitness')
plt.savefig('Figure-5b.png', dpi=300)
plt.show()

# %% [markdown]
### Channel Capacity Depending on the folding state

# %% [code]
# Initial Condition of simulations by varying phi_fold
import numpy as np
from pysb.simulator import ScipyOdeSimulator
import matplotlib.pyplot as plt

def update_initial_conditions(model, initial_conditions):
    for monomer, value in initial_conditions.items():
        model.parameters[f"{monomer}_0"].value = value

def update_folding_parameters(model, folding_conditions):
    """
    Update folding-related parameters.
    Expects a dictionary with keys matching parameter names in the model,
    for example: {'phi_fold': value, 'Delta_G_fold': value}
    """
    for param, value in folding_conditions.items():
        model.parameters[param].value = value

# Time span for simulation (e.g., 4 days in seconds)
tspan = np.linspace(0, 345600, 115200)
results = []
# Define ranges for phi_fold (folding state) and Delta_G_fold.
# Here, phi_fold varies from 0 to 1 (11 intervals),
# and Delta_G_fold varies over an example range (e.g., 150 to 200 kcal/mol) in 11 intervals.
phi_values = np.linspace(0, 1, 10)          # φ: 0, 0.1, 0.2, ..., 1.0
# Fixed initial conditions for species that are not varied:
initial_conditions = {
    'eif2alpha': 2244,  # constant value (adjust as needed)
    'actin': 2400,      # constant value
    'GADD34': 40,       # fixed GADD34 concentration
    'PP1': 800          # fixed PP1 concentration
}
update_initial_conditions(model, initial_conditions)

# Loop over phi_fold and Delta_G_fold values.
for i, phi_value in enumerate(phi_values):
    folding_conditions = {
    'phi_fold': phi_value,
    'Delta_G_fold': 170,
    'Delta_G_actin': 15.7,
    'Delta_G_PP1_trimer':40.661,
    'Delta_G_PP1_dimer': 40.661
            
            }
    update_folding_parameters(model, folding_conditions)
        
    # Create a new simulator instance (using LSODA with specified tolerances)
    simulator = ScipyOdeSimulator(
                 model,
                tspan=tspan,
                integrator_options={'rtol': 1e-12, 'atol': 1e-12}
            )
    sim_result = simulator.run()
    results.append(sim_result.all)

# %% [code]
import pandas as pd
import matplotlib.pyplot as plt
# Extract parameters from the model
data = [(p.name, p.value) for p in model.parameters]
df = pd.DataFrame(data, columns=['Parameter', 'Value'])

# Set up the matplotlib figure
fig, ax = plt.subplots(figsize=(6, 3))
ax.axis('tight')
ax.axis('off')

# Create the table and center-align the text
table = ax.table(cellText=df.values,
                 colLabels=df.columns,
                 cellLoc='center',
                 loc='center')

# Improve table aesthetics
table.auto_set_font_size(False)
table.set_fontsize(12)
table.scale(1.2, 1.2)

# Bold the header cells (column names)
for key, cell in table.get_celld().items():
    # Header cells are in the first row (row index 0)
    if key[0] == 0:
        cell.get_text().set_fontweight('bold')

# Optionally, add a title
#plt.title("PySB Model Parameters", fontsize=14)

# Save the figure with bbox_inches='tight' to capture the full table
plt.savefig("pysb_parameters_fromMD.png", bbox_inches='tight')
plt.show()

# %% [code]
import matplotlib.pyplot as plt
import matplotlib.ticker as ticker

# Your time span in seconds:
tspan = np.linspace(0, 345600, 115200)

# Define tick positions every 12 hours (43,200 s)
tick_positions = np.arange(0, 345600 + 1, 43200)  # +1 to include 345600

# Create corresponding tick labels in hours by dividing seconds by 3600:
tick_labels = tick_positions / 3600


# Assume cmap, tspan, results, and kinetic_conditions_list are already defined
# For example:
cmap = plt.get_cmap('viridis')
# tspan = ...   (your time array)
# results = run__kinetic_simulations(model, kinetic_conditions_list, tspan)

# Create a figure with 4 rows and 2 columns (8 subplots)
fig, axs = plt.subplots(4, 2, figsize=(16, 10))
axs = axs.flatten()  # Flatten to index subplots as a 1D array

# Subplot 0: GADD34 Over Time (using the 'ppp1r15a' data)
for idx, result in enumerate(results):
    phi_fold = phi_values[idx]
    color = cmap(phi_fold)
    axs[0].plot(tspan, result['ppp1r15a'], label=f"phi_fold = {phi_fold}", color=color)
axs[0].set_ylabel('GADD34 [mM]')
axs[0].set_xticklabels([])
axs[0].grid(True, axis='x', which='major', linestyle='--', color='gray', alpha=0.7)

# Subplot 1: GADD34 actin Dimer Over Time
for idx, result in enumerate(results):
    phi_fold = phi_values[idx]
    color = cmap(phi_fold)
    axs[1].plot(tspan, result['GADD34_actin'], label=f"phi_fold = {phi_fold}", color=color)
axs[1].set_ylabel('actin:GADD34 [mM]')
axs[1].set_xticklabels([])
axs[1].grid(True, axis='x', which='major', linestyle='--', color='gray', alpha=0.7)
#axs[1].set_title('GADD34 actin Dimer Over Time')

# Subplot 2: PP1 trimer Over Time
for idx, result in enumerate(results):
    phi_fold = phi_values[idx]
    color = cmap(phi_fold)
    axs[2].plot(tspan, result['PP1_trimer_folded'], label=f"phi_fold = {phi_fold}", color=color)
axs[2].set_ylabel('actin:GADD34:PP1 [mM]')
axs[2].set_xticklabels([])
axs[2].grid(True, axis='x', which='major', linestyle='--', color='gray', alpha=0.7)


# Subplot 3: Holoenzyme Formation Over Time
for idx, result in enumerate(results):
    phi_fold = phi_values[idx]
    color = cmap(phi_fold)
    axs[3].plot(tspan, result['holoenzyme'], label=f"$\phi$ = {phi_fold}", color=color)
axs[3].set_ylabel('actin:GADD34:PP1:eif2$\\alpha$[mM]')
axs[3].set_xticklabels([])
axs[3].grid(True, axis='x', which='major', linestyle='--', color='gray', alpha=0.7)
axs[3].legend(bbox_to_anchor=(1, 1))


# Subplot 4: GADD34 PP1 Dimer Over Time
for idx, result in enumerate(results):
    phi_fold = phi_values[idx]
    color = cmap(phi_fold) 
    axs[4].plot(tspan, result['GADD34_PP1'], label=f"$\phi$ = {phi_fold}", color=color) 
axs[4].set_ylabel('GADD34:PP1 [mM]')
axs[4].set_xticklabels([])
axs[4].grid(True, axis='x', which='major', linestyle='--', color='gray', alpha=0.7)

# Subplot 5: PP1 trimer Over Time
for idx, result in enumerate(results):
    phi_fold = phi_values[idx]
    color = cmap(phi_fold)
    axs[5].plot(tspan, result['PP1_trimer_formed'], label=f"phi_fold = {phi_fold}", color=color)
axs[5].set_ylabel('GADD34:PP1:eif2$\\alpha$ [mM]')
axs[5].set_xticklabels([])
axs[5].grid(True, axis='x', which='major', linestyle='--', color='gray', alpha=0.7)


# Subplot 6: eif2alpha-phosphorylated Over Time
for idx, result in enumerate(results):
    phi_fold = phi_values[idx]
    color = cmap(phi_fold)
    axs[6].plot(tspan, result['phospho_eif2alpha'], label=f"phi_fold = {phi_fold}", color=color)
axs[6].set_xlabel('Time [hours]')
axs[6].set_ylabel('phospho-eif2$\\alpha$ [mM]')
axs[6].set_xticks(tick_positions)
axs[6].set_xticklabels(tick_labels)
axs[6].grid(True, axis='x', which='major', linestyle='--', color='gray', alpha=0.7)

# Subplot 7: eif2alpha-dephosphorylated Over Time
for idx, result in enumerate(results):
    phi_fold = phi_values[idx]
    color = cmap(phi_fold)
    axs[7].plot(tspan, result['dephospho_eif2alpha'], label=f"phi_fold = {phi_fold}", color=color)
axs[7].set_xlabel('Time [hours]')
axs[7].set_ylabel("eif2$\\alpha$ [mM]")
axs[7].set_xticks(tick_positions)
axs[7].set_xticklabels(tick_labels)
axs[7].grid(True, axis='x', which='major', linestyle='--', color='gray', alpha=0.7)


plt.tight_layout()
plt.savefig('Figure-S4.png', dpi=300)
plt.show()

# %% [markdown]
### Channel Capacity

# %% [markdown]
### Monitoring the channel capacity across two channels, defined by the following reactions:
- Channel 1: GADD34 -> GADD34:actin -> PP1:GADD34:actin -> PP1:GADD34:actin:p-eIF2alpha
- Channel 2: GADD34 -> GADD34:PP1 -> GADD34:PP1:eif2alpha
- Consider comparing the two signal given the transcription factor activity (ATF4)

# %% [code]
import numpy as np

def normalize_array(arr, method='minmax'):
    """
    Normalize a NumPy array using the specified method.
    
    Parameters:
        arr (array-like): The input array to normalize.
        method (str): The normalization method. Options:
                      - 'minmax': Scales the array to the range [0, 1].
                      - 'zscore': Standardizes the array to have mean 0 and std 1.
                      
    Returns:
        np.ndarray: The normalized array.
    """
    arr = np.array(arr, dtype=np.float64)
    
    if method == 'minmax':
        # Compute minimum and maximum values, ignoring NaNs.
        min_val = np.nanmin(arr)
        max_val = np.nanmax(arr)
        range_val = max_val - min_val
        if range_val == 0:
            return np.zeros_like(arr)
        return (arr - min_val) / range_val
    elif method == 'zscore':
        # Compute mean and standard deviation, ignoring NaNs.
        mean_val = np.nanmean(arr)
        std_val = np.nanstd(arr)
        if std_val == 0:
            return np.zeros_like(arr)
        return (arr - mean_val) / std_val
    else:
        raise ValueError("Normalization method not recognized. Use 'minmax' or 'zscore'.")

# %% [code]
import numpy as np
import pyinform as pinform
import matplotlib.pyplot as plt

# --- Data Collection ---
# Assuming 'results' is a list of simulation outputs and 'phi_values' contains the corresponding φ_fold values.
fold_values = []
mutual_information_values = []
mutual_information_values_1 = []
mutual_information_values_2 = []
mutual_information_values_a = []
mutual_information_values_b = []


for idx, result in enumerate(results):
    # Extract the folding value for this simulation
    phi_fold = phi_values[idx]
    fold_values.append(phi_fold)
    
    # Select observables for the analysis
    u = result['ppp1r15a']          # Input observable (GADD34(..., s='u'))
    y = result['GADD34_folded']
    x = result['GADD34_PP1']
    z = result['PP1_trimer_folded']
    v = result['PP1_trimer_formed']
    w = result['holoenzyme']
    # Output observable (actin(...) % GADD34(..., s='u'))
    
    # Normalize signals (assuming normalize_array is defined elsewhere)
    u_normalized = normalize_array(u, method='minmax')
    y_normalized = normalize_array(y, method='minmax')
    x_normalized = normalize_array(x, method='minmax')
    z_normalized = normalize_array(z, method='minmax')
    v_normalized = normalize_array(v, method='minmax')
    w_normalized = normalize_array(w, method='minmax')
    
    
    # Compute mutual information between the normalized signals
    mi = pinform.mutual_info(u_normalized, y_normalized)
    mutual_information_values.append(mi)
    mi_1 = pinform.mutual_info(y_normalized, z_normalized)
    mutual_information_values_1.append(mi_1)
    mi_2 = pinform.mutual_info(z_normalized, w_normalized)
    mutual_information_values_2.append(mi_2)
    mi_a = pinform.mutual_info(u_normalized, x_normalized)
    mutual_information_values_a.append(mi_a)
    mi_b = pinform.mutual_info(x_normalized, v_normalized)
    mutual_information_values_b.append(mi_b)
    

# Compute the overall channel capacity as the maximum mutual information value
channel_capacity = max(mutual_information_values)
#print("Computed Channel Capacity R1:", channel_capacity)
channel_capacity_1 = max(mutual_information_values_1)
#print("Computed Channel Capacity R2:", channel_capacity_1)
channel_capacity_2 = max(mutual_information_values_2)
#print("Computed Channel Capacity R3:", channel_capacity_2)
channel_capacity_a = max(mutual_information_values_a)
#print("Computed Channel Capacity R4:", channel_capacity_a)
channel_capacity_b = max(mutual_information_values_b)
#print("Computed Channel Capacity R5:", channel_capacity_b)


# --- Figure 2: Violin Plot of Mutual Information Distribution ---
plt.figure(figsize=(8, 6))

# Combine all mutual information data into a list
data = [
    mutual_information_values, 
    mutual_information_values_1, 
    mutual_information_values_2, 
    mutual_information_values_a, 
    mutual_information_values_b
]
# Define positions for each violin plot
positions = [1, 2, 3, 4, 5]

# Create the violin plot for each data set at specified positions
plt.violinplot(data, positions=positions, showmeans=False, showextrema=False, showmedians=True)

# Optionally add horizontal lines for channel capacities from each channel if desired
plt.axhline(max(mutual_information_values), color='r', linestyle='--', label=f'Channel Capacity GADD34 <-> actin:GADD34 = {max(mutual_information_values):.4f}')
plt.axhline(max(mutual_information_values_1), color='g', linestyle='--', label=f'Channel Capacity actin:GADD34 <-> actin:GADD34:PP1 = {max(mutual_information_values_1):.4f}')
plt.axhline(max(mutual_information_values_2), color='b', linestyle='--', label=f'Channel Capacity actin:GADD34:PP1 <-> actin:GADD34:PP1:eif2$\\alpha$ = {max(mutual_information_values_2):.4f}')
plt.axhline(max(mutual_information_values_a), color='m', linestyle='--', label=f'Channel Capacity GADD34 <-> GADD34:PP1 = {max(mutual_information_values_a):.4f}')
plt.axhline(max(mutual_information_values_b), color='c', linestyle='--', label=f'Channel Capacity  GADD34:PP1 <-> GADD34:PP1:eif2$\\alpha$ = {max(mutual_information_values_b):.4f}')

plt.ylabel('Mutual Information', fontsize=16)
# Update x-tick labels to correspond to each channel
plt.xticks(positions, ['GADD34 <-> actin:GADD34', 'actin:GADD34 <-> actin:GADD34:PP1', 'actin:GADD34:PP1 <-> actin:GADD34:PP1:eif2$\\alpha$', 'GADD34 <-> GADD34:PP1', 'GADD34:PP1 <-> GADD34:PP1:eif2$\\alpha$'], rotation=45)
#plt.title('Violin Plot of Mutual Information Distribution', fontsize=18)
plt.legend(fontsize=12, bbox_to_anchor=(1.05, 1), loc='upper left')
#plt.grid(True, linestyle='--', alpha=0.7)
plt.savefig('Figure-6a.png', dpi=300)
plt.show()


# %% [markdown]
## Channel Capacity from Optimal State from phase space

# %% [code]
# Initial Condition of simulations by varying phi_fold
def update_initial_conditions(model, initial_conditions):
    for monomer, value in initial_conditions.items():
        model.parameters[f"{monomer}_0"].value = value

def update_folding_parameters(model, folding_conditions):
    """
    Update folding-related parameters.
    Expects a dictionary with keys matching parameter names in the model,
    for example: {'phi_fold': value, 'Delta_G_fold': value}
    """
    for param, value in folding_conditions.items():
        model.parameters[param].value = value

# Time span for simulation (e.g., 4 days in seconds)
tspan = np.linspace(0, 345600, 115200)
results = []
# Define ranges for phi_fold (folding state) and Delta_G_fold.
# Here, phi_fold varies from 0 to 1 (11 intervals),
# and Delta_G_fold varies over an example range (e.g., 150 to 200 kcal/mol) in 11 intervals.
phi_values = np.linspace(0, 1, 10)          # φ: 0, 0.1, 0.2, ..., 1.0
# Fixed initial conditions for species that are not varied:
initial_conditions = {
    'eif2alpha': 2244,  # constant value (adjust as needed)
    'actin': 2400,      # constant value
    'GADD34': 40,       # fixed GADD34 concentration
    'PP1': 800          # fixed PP1 concentration
}
update_initial_conditions(model, initial_conditions)

# Loop over phi_fold and Delta_G_fold values.
for i, phi_value in enumerate(phi_values):
    folding_conditions = {
    'phi_fold': phi_value,
    'Delta_G_fold': 20,
    'Delta_G_actin': 3.3,
    'Delta_G_PP1_trimer':38.8,
    'Delta_G_PP1_dimer': 38.8
            
            }
    update_folding_parameters(model, folding_conditions)
        
    # Create a new simulator instance (using LSODA with specified tolerances)
    simulator = ScipyOdeSimulator(
                 model,
                tspan=tspan,
                integrator_options={'rtol': 1e-12, 'atol': 1e-12}
            )
    sim_result = simulator.run()
    results.append(sim_result.all)

# %% [code]
import pandas as pd
import matplotlib.pyplot as plt
# Extract parameters from the model
data = [(p.name, p.value) for p in model.parameters]
df = pd.DataFrame(data, columns=['Parameter', 'Value'])

# Set up the matplotlib figure
fig, ax = plt.subplots(figsize=(6, 3))
ax.axis('tight')
ax.axis('off')

# Create the table and center-align the text
table = ax.table(cellText=df.values,
                 colLabels=df.columns,
                 cellLoc='center',
                 loc='center')

# Improve table aesthetics
table.auto_set_font_size(False)
table.set_fontsize(12)
table.scale(1.2, 1.2)

# Bold the header cells (column names)
for key, cell in table.get_celld().items():
    # Header cells are in the first row (row index 0)
    if key[0] == 0:
        cell.get_text().set_fontweight('bold')

# Optionally, add a title
#plt.title("PySB Model Parameters", fontsize=14)

# Save the figure with bbox_inches='tight' to capture the full table
plt.savefig("pysb_parameters_fromOptimalPhase.png", bbox_inches='tight')
plt.show()

# %% [code]
import matplotlib.pyplot as plt
import matplotlib.ticker as ticker

# Your time span in seconds:
tspan = np.linspace(0, 345600, 115200)

# Define tick positions every 12 hours (43,200 s)
tick_positions = np.arange(0, 345600 + 1, 43200)  # +1 to include 345600

# Create corresponding tick labels in hours by dividing seconds by 3600:
tick_labels = tick_positions / 3600


# Assume cmap, tspan, results, and kinetic_conditions_list are already defined
# For example:
# cmap = plt.get_cmap('viridis')
# tspan = ...   (your time array)
# results = run__kinetic_simulations(model, kinetic_conditions_list, tspan)

# Create a figure with 4 rows and 2 columns (8 subplots)
fig, axs = plt.subplots(4, 2, figsize=(16, 10))
axs = axs.flatten()  # Flatten to index subplots as a 1D array

# Subplot 0: GADD34 Over Time (using the 'ppp1r15a' data)
for idx, result in enumerate(results):
    phi_fold = phi_values[idx]
    color = cmap(phi_fold)
    axs[0].plot(tspan, result['ppp1r15a'], label=f"phi_fold = {phi_fold}", color=color)
axs[0].set_ylabel('GADD34 [mM]')
axs[0].set_xticklabels([])
axs[0].grid(True, axis='x', which='major', linestyle='--', color='gray', alpha=0.7)

# Subplot 1: GADD34 actin Dimer Over Time
for idx, result in enumerate(results):
    phi_fold = phi_values[idx]
    color = cmap(phi_fold)
    axs[1].plot(tspan, result['GADD34_actin'], label=f"phi_fold = {phi_fold}", color=color)
axs[1].set_ylabel('actin:GADD34 [mM]')
axs[1].set_xticklabels([])
axs[1].grid(True, axis='x', which='major', linestyle='--', color='gray', alpha=0.7)
#axs[1].set_title('GADD34 actin Dimer Over Time')

# Subplot 2: PP1 trimer Over Time
for idx, result in enumerate(results):
    phi_fold = phi_values[idx]
    color = cmap(phi_fold)
    axs[2].plot(tspan, result['PP1_trimer_folded'], label=f"phi_fold = {phi_fold}", color=color)
axs[2].set_ylabel('actin:GADD34:PP1 [mM]')
axs[2].set_xticklabels([])
axs[2].grid(True, axis='x', which='major', linestyle='--', color='gray', alpha=0.7)


# Subplot 3: Holoenzyme Formation Over Time
for idx, result in enumerate(results):
    phi_fold = phi_values[idx]
    color = cmap(phi_fold)
    axs[3].plot(tspan, result['holoenzyme'], label=f"$\phi$ = {phi_fold}", color=color)
axs[3].set_ylabel('actin:GADD34:PP1:eif2$\\alpha$[mM]')
axs[3].set_xticklabels([])
axs[3].grid(True, axis='x', which='major', linestyle='--', color='gray', alpha=0.7)
axs[3].legend(bbox_to_anchor=(1, 1))


# Subplot 4: GADD34 PP1 Dimer Over Time
for idx, result in enumerate(results):
    phi_fold = phi_values[idx]
    color = cmap(phi_fold) 
    axs[4].plot(tspan, result['GADD34_PP1'], label=f"$\phi$ = {phi_fold}", color=color) 
axs[4].set_ylabel('GADD34:PP1 [mM]')
axs[4].set_xticklabels([])
axs[4].grid(True, axis='x', which='major', linestyle='--', color='gray', alpha=0.7)

# Subplot 5: PP1 trimer Over Time
for idx, result in enumerate(results):
    phi_fold = phi_values[idx]
    color = cmap(phi_fold)
    axs[5].plot(tspan, result['PP1_trimer_formed'], label=f"phi_fold = {phi_fold}", color=color)
axs[5].set_ylabel('GADD34:PP1:eif2$\\alpha$ [mM]')
axs[5].set_xticklabels([])
axs[5].grid(True, axis='x', which='major', linestyle='--', color='gray', alpha=0.7)


# Subplot 6: eif2alpha-phosphorylated Over Time
for idx, result in enumerate(results):
    phi_fold = phi_values[idx]
    color = cmap(phi_fold)
    axs[6].plot(tspan, result['phospho_eif2alpha'], label=f"phi_fold = {phi_fold}", color=color)
axs[6].set_xlabel('Time [hours]')
axs[6].set_ylabel('phospho-eif2$\\alpha$ [mM]')
axs[6].set_xticks(tick_positions)
axs[6].set_xticklabels(tick_labels)
axs[6].grid(True, axis='x', which='major', linestyle='--', color='gray', alpha=0.7)

# Subplot 7: eif2alpha-dephosphorylated Over Time
for idx, result in enumerate(results):
    phi_fold = phi_values[idx]
    color = cmap(phi_fold)
    axs[7].plot(tspan, result['dephospho_eif2alpha'], label=f"phi_fold = {phi_fold}", color=color)
axs[7].set_xlabel('Time [hours]')
axs[7].set_ylabel("eif2$\\alpha$ [mM]")
axs[7].set_xticks(tick_positions)
axs[7].set_xticklabels(tick_labels)
axs[7].grid(True, axis='x', which='major', linestyle='--', color='gray', alpha=0.7)


plt.tight_layout()
plt.savefig('Figure-S5.png', dpi=300)
plt.show()

# %% [code]
#Adjust size of the table
from util_display import format_species_reactions, display_table
import pandas as pd
# Prevent pandas from truncating long LaTeX expressions when rendering.
pd.options.display.max_colwidth=None

#obtain dataframe with math latex expression visualization for species and reactions
(speciesdisp, reactionsdisp)=format_species_reactions(model);
#display species with the right style
display_table(speciesdisp, caption='SPECIES')

#display reactions in the same order as Reaction List S3 in original publication 
#reaction_order=[0,1,2,4,5,5]
display_table(reactionsdisp, caption='REACTIONS')
#display_table(reactionsdisp.iloc[reaction_order], caption='REACTIONS')

# %% [code]
def safe_series(ts, bins=100, verbose=True):
    """
    Convert a time series to discrete nonnegative integer states.
    
    1. Shifts the series if any negative values are present.
    2. Discretizes the (shifted) series into a specified number of bins.
    
    Parameters:
        ts (array-like): The input time series.
        bins (int): Number of discrete bins to use.
        verbose (bool): If True, prints out information about the adjustments.
    
    Returns:
        np.ndarray: Discretized time series with integer values.
    """
    ts = np.asarray(ts)
    
    # Shift series if negative values exist
    if ts.min() < 0:
        shift = -ts.min()
        if verbose:
            print(f"Shifting series by {shift} to ensure nonnegative values.")
        ts = ts + shift
    
    # If the series is constant, return an array of zeros
    if ts.max() == ts.min():
        if verbose:
            print("Series is constant; returning zeros.")
        return np.zeros_like(ts, dtype=int)
    
    # Discretize the series into 'bins' number of states
    # The states will be in the range [0, bins - 1]
    ts_scaled = np.floor((ts - ts.min()) / (ts.max() - ts.min()) * (bins - 1)).astype(int)
    return ts_scaled

# %% [code]
import numpy as np
import pyinform as pinform
import matplotlib.pyplot as plt

# --- Data Collection ---
# Assuming 'results' is a list of simulation outputs and 'phi_values' contains the corresponding φ_fold values.
fold_values = []
mutual_information_values = []
mutual_information_values_1 = []
mutual_information_values_2 = []
mutual_information_values_a = []
mutual_information_values_b = []


for idx, result in enumerate(results):
    # Extract the folding value for this simulation
    phi_fold = phi_values[idx]
    fold_values.append(phi_fold)
    
    # Select observables for the analysis
    u = result['ppp1r15a']          # Input observable (GADD34(..., s='u'))
    y = result['GADD34_folded']
    x = result['GADD34_PP1']
    z = result['PP1_trimer_folded']
    v = result['PP1_trimer_formed']
    w = result['holoenzyme']
    # Output observable (actin(...) % GADD34(..., s='u'))
    
    # Normalize signals (assuming normalize_array is defined elsewhere)
    u_normalized = normalize_array(u, method='minmax')
    y_normalized = normalize_array(y, method='minmax')
    x_normalized = normalize_array(x, method='minmax')
    z_normalized = normalize_array(z, method='minmax')
    v_normalized = normalize_array(v, method='minmax')
    w_normalized = normalize_array(w, method='minmax')
    
    # Ensure all arrays have nonnegative values
    u_nonneg = safe_series(u)
    y_nonneg = safe_series(y)
    z_nonneg = safe_series(z)
    w_nonneg = safe_series(w)
    x_nonneg = safe_series(x)
    v_nonneg = safe_series(v)
    
    
    # Compute mutual information between the normalized signals
    mi = pinform.mutual_info(u_nonneg, y_nonneg)
    mutual_information_values.append(mi)
    mi_1 = pinform.mutual_info(y_nonneg, z_nonneg)
    mutual_information_values_1.append(mi_1)
    mi_2 = pinform.mutual_info(z_nonneg, w_nonneg)
    mutual_information_values_2.append(mi_2)
    mi_a = pinform.mutual_info(u_nonneg, x_nonneg)
    mutual_information_values_a.append(mi_a)
    mi_b = pinform.mutual_info(x_nonneg, v_nonneg)
    mutual_information_values_b.append(mi_b)
    

# Compute the overall channel capacity as the maximum mutual information value
channel_capacity = max(mutual_information_values)
#print("Computed Channel Capacity R1:", channel_capacity)
channel_capacity_1 = max(mutual_information_values_1)
#print("Computed Channel Capacity R2:", channel_capacity_1)
channel_capacity_2 = max(mutual_information_values_2)
#print("Computed Channel Capacity R3:", channel_capacity_2)
channel_capacity_a = max(mutual_information_values_a)
#print("Computed Channel Capacity R4:", channel_capacity_a)
channel_capacity_b = max(mutual_information_values_b)
#print("Computed Channel Capacity R5:", channel_capacity_b)


# --- Figure 2: Violin Plot of Mutual Information Distribution ---
plt.figure(figsize=(8, 6))

# Combine all mutual information data into a list
data = [
    mutual_information_values, 
    mutual_information_values_1, 
    mutual_information_values_2, 
    mutual_information_values_a, 
    mutual_information_values_b
]
# Define positions for each violin plot
positions = [1, 2, 3, 4, 5]

# Create the violin plot for each data set at specified positions
plt.violinplot(data, positions=positions, showmeans=False, showextrema=False, showmedians=True)

# Optionally add horizontal lines for channel capacities from each channel if desired
plt.axhline(max(mutual_information_values), color='r', linestyle='--', label=f'Channel Capacity GADD34 <-> actin:GADD34 = {max(mutual_information_values):.4f}')
plt.axhline(max(mutual_information_values_1), color='g', linestyle='--', label=f'Channel Capacity actin:GADD34 <-> actin:GADD34:PP1 = {max(mutual_information_values_1):.4f}')
plt.axhline(max(mutual_information_values_2), color='b', linestyle='--', label=f'Channel Capacity actin:GADD34:PP1 <-> actin:GADD34:PP1:eif2$\\alpha$ = {max(mutual_information_values_2):.4f}')
plt.axhline(max(mutual_information_values_a), color='m', linestyle='--', label=f'Channel Capacity GADD34 <-> GADD34:PP1 = {max(mutual_information_values_a):.4f}')
plt.axhline(max(mutual_information_values_b), color='c', linestyle='--', label=f'Channel Capacity  GADD34:PP1 <-> GADD34:PP1:eif2$\\alpha$ = {max(mutual_information_values_b):.4f}')

plt.ylabel('Mutual Information', fontsize=16)
# Update x-tick labels to correspond to each channel
plt.xticks(positions, ['GADD34 <-> actin:GADD34', 'actin:GADD34 <-> actin:GADD34:PP1', 'actin:GADD34:PP1 <-> actin:GADD34:PP1:eif2$\\alpha$', 'GADD34 <-> GADD34:PP1', 'GADD34:PP1 <-> GADD34:PP1:eif2$\\alpha$'], rotation=45)
#plt.title('Violin Plot of Mutual Information Distribution', fontsize=18)
plt.legend(fontsize=12, bbox_to_anchor=(1.05, 1), loc='upper left')
#plt.grid(True, linestyle='--', alpha=0.7)
plt.savefig('Figure-6b.png', dpi=300)
plt.show()

# %% [code]


# %% [code]
```
