## Supplementary material for "Folding-Driven Control of the Functional PP1 Complex via Multiscale Modeling": Supporting_Information.pdf

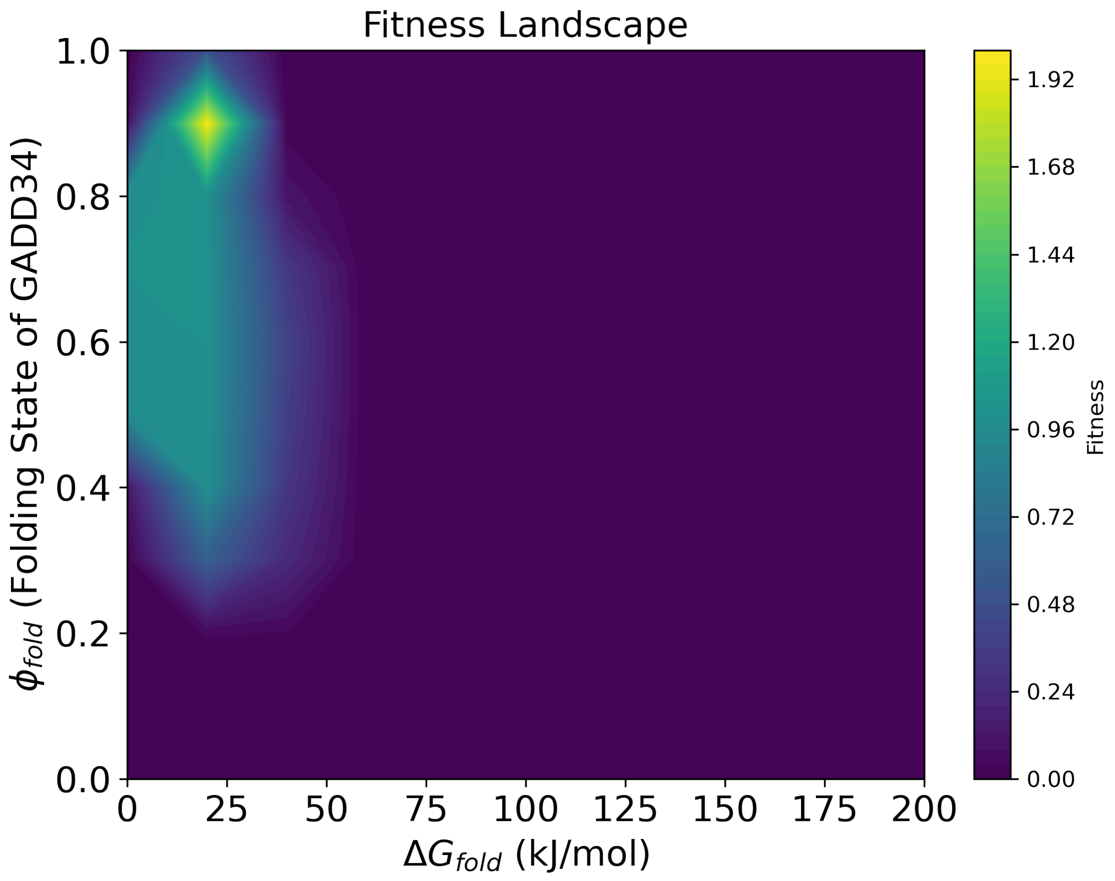


**Figure S1.** Two-dimensional heatmap illustrating the efficiency of PP1 complex enzymatic activity (color scale) as a function of $\Delta G_{fold}$ (x-axis) and ϕ, unitless (y-axis) of GADD34. The maximum fitness is reached for $\Delta G_{fold}$ equal to 20 kJ/mol and ϕ equal to 0.9.

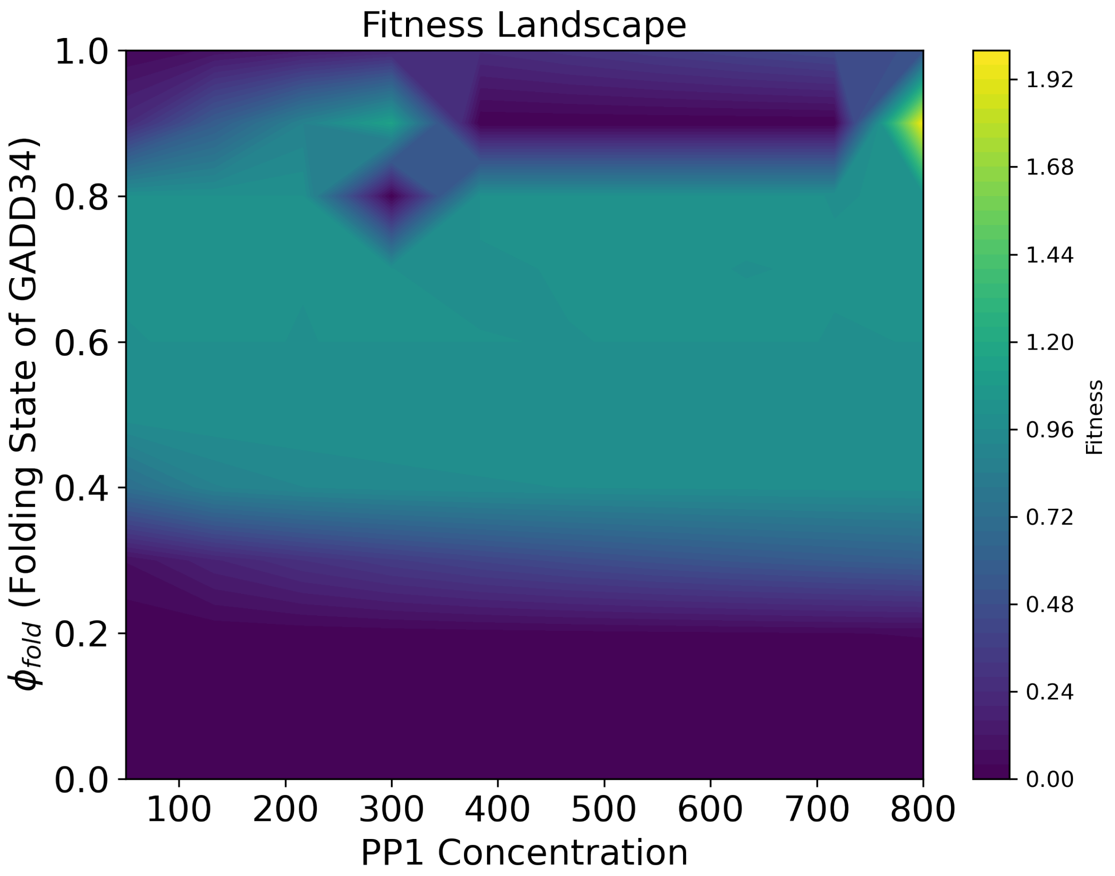


**Figure S2.** Two-dimensional heatmap illustrating the efficiency of PP1 complex enzymatic activity (color scale) as a function of PP1 concentration (x-axis) and **ϕ, unitless,** (y-axis) of GADD34. The maximum fitness is reached for the PP1 concentration of 800 mM and **ϕ** equal to 0.9.

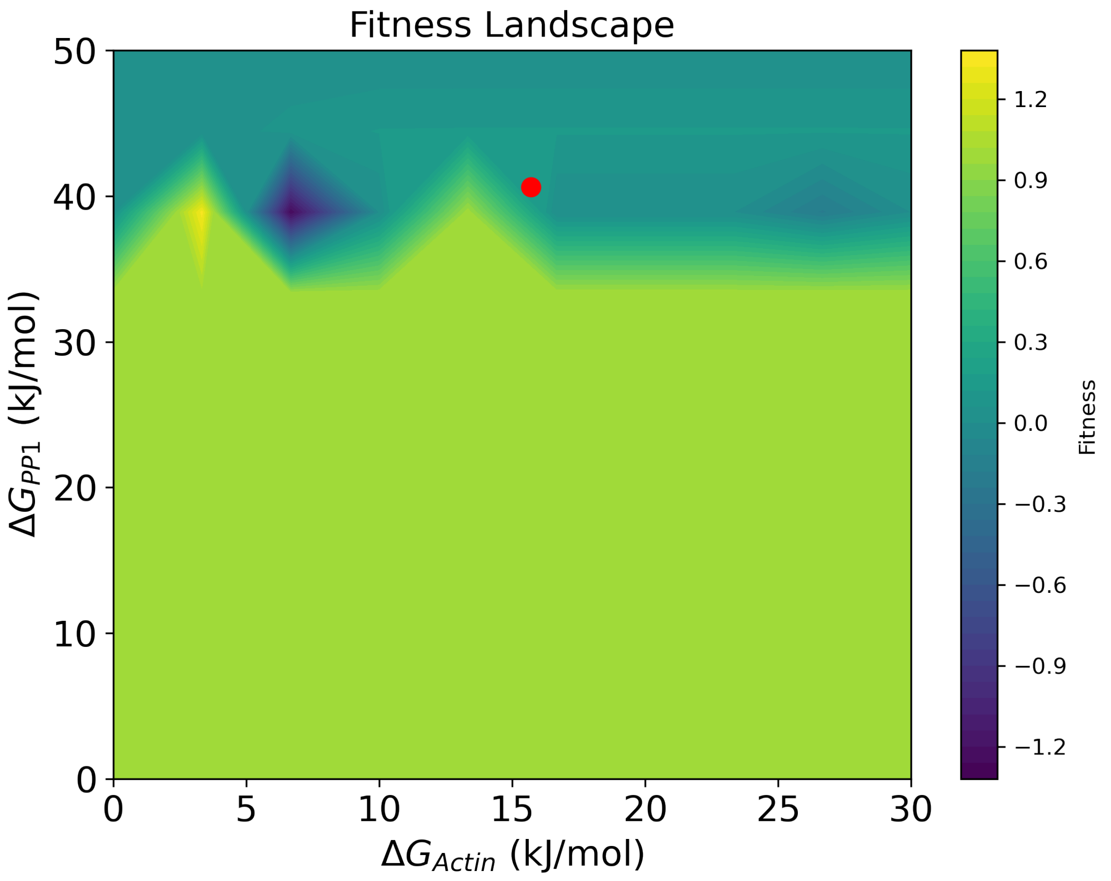


**Figure S3.** Two-dimensional heatmap illustrating the efficiency of PP1 complex enzymatic activity, PP1 concentration equal to 800 mM, ϕ equal to 0.9, $\Delta G_{fold}$ equal to 20 kJ/mol, (color scale) as a function of $\Delta G_{actin}$ (x-axis) and $\Delta G_{PP1}$ (y-axis), relative to the binding of GADD34 to actin and PP1, respectively. The maximum fitness is reached for $\Delta G_{actin}$ and $\Delta G_{PP1}$ equal to 3.3 kJ/mol and 38.8 kJ/mol, respectively. The red dot points to the conditions sampled through umbrella sampling MD simulations.

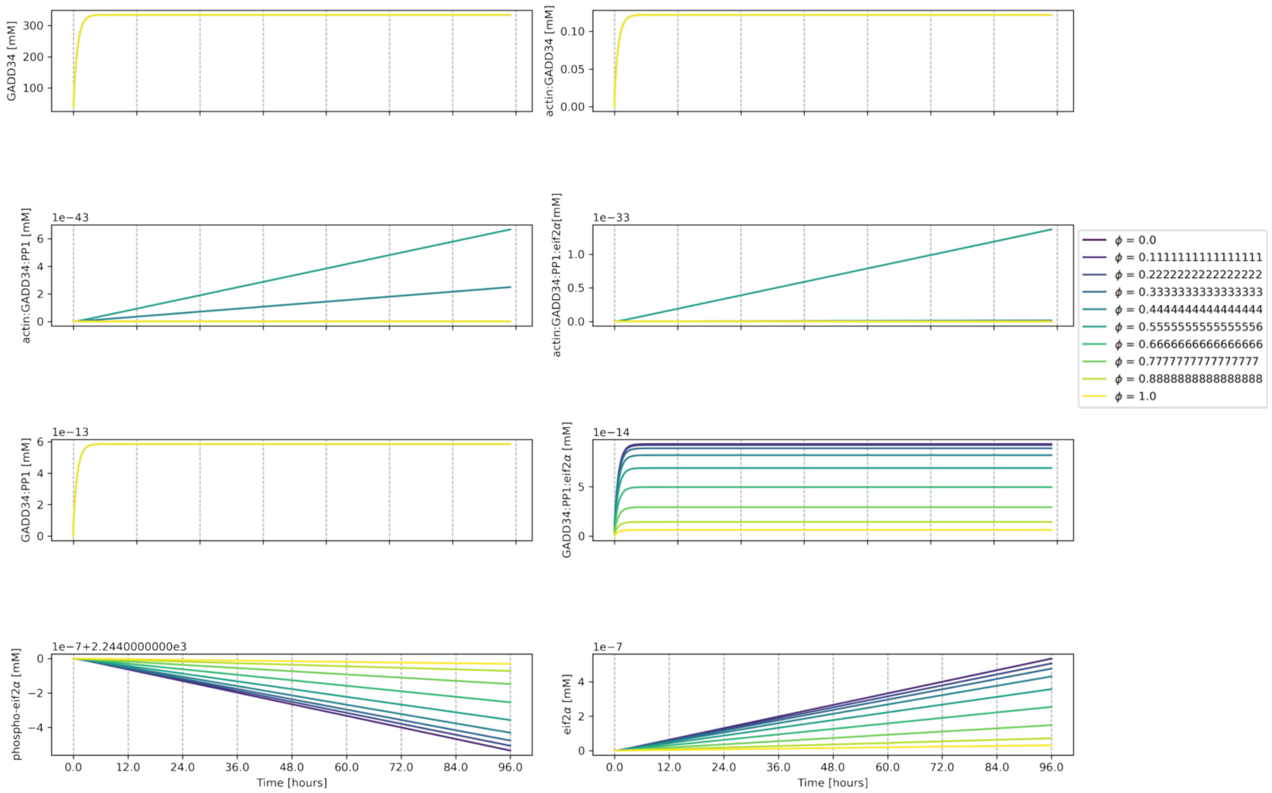


**Figure S4.** The time course of intermediate dimers that concur to the formation of PP1 holoenzyme and dephosphorylation of phosphorylated-eif2$\alpha$, as a function of **ϕ,** given $\Delta G_{actin}$ equally to 15.7 kJ/mol, $\Delta G_{PP1}$ equal to 40.661 kJ/mol, and $\Delta G_{fold}$ equal to 170 kJ/mol.

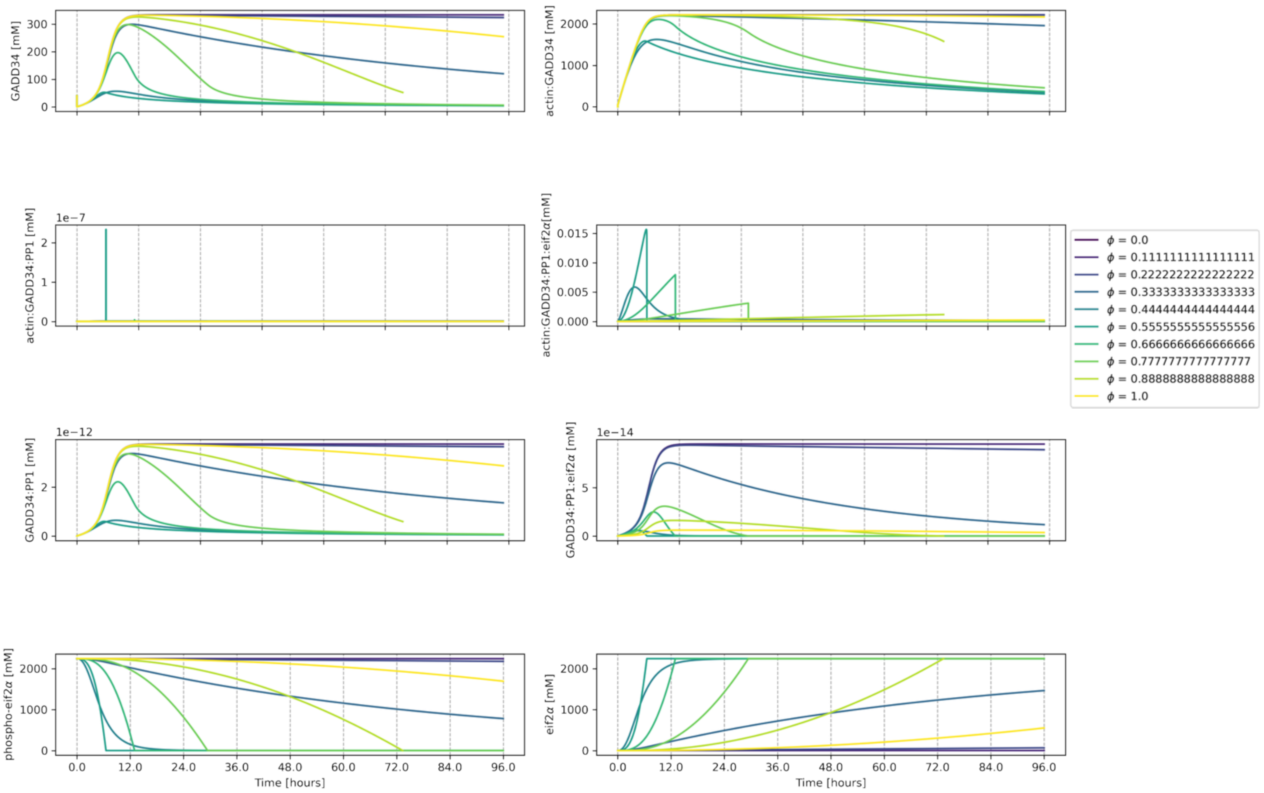


**Figure S5.** The time course of intermediate dimers that concur to the formation of PP1 holoenzyme and dephosphorylation of phosphor-eif2$\alpha$, as a function of **ϕ,** given $\Delta G_{actin}$ equally to 3.3 kJ/mol, $\Delta G_{PP1}$ equal to 38.8 kJ/mol, and $\Delta G_{fold}$ equal to 20 kJ/mol.

**Table S1.** Parameters used in the PySB model for the PP1–GADD34–eIF2α–actin system. Two values are shown: the "Energy Constrained State" and the "Optimal State," highlighting initial monomer concentrations, key kinetic rates, and thermodynamic constraints. The parameter φ_fold (phi_fold) governs GADD34's conformational dynamics, influencing complex assembly and function.

| Parameter | Value – Energy Constrained State | Value – Optimal State |
|---|---|---|
| **Monomers** | | |
| GADD34_0 [mM] | 40.0 | 40.0 |
| eif2alpha_0 [mM] | 2244.0 | 2244.0 |
| actin_0 [mM] | 2400.0 | 2400.0 |
| PP1_0 [mM] | 800 | 800 |
| **Kinetic Constants** | | |
| phi_fold | 0 to 1 | 0 to 1 |
| kdeg_GADD34 [$s^{-1}$] | 0.000309 | 0.000309 |
| ksyn_GADD34[$s^{-1}$] | 4.6e-5 | 4.6e-5 |
| kcat_PP1[$s^{-1}$] | 2.23e6 | 2.23e6 |
| **Energy Constraints** | | |
| $\Delta$G_actin [kJ/mol] | 15.7 | 3.3 |
| $\Delta$G_PP1 [kJ/mol] | 40.66 | 38.8 |
| $\Delta$G_eif2$\alpha$_PP1 [kJ/mol] | 50.189 | 50.189 |
| $\Delta$G_eif2 $\alpha$ | 10.0 | 10.0 |
